# The effect of estradiol during the early stages of osteoclast differentiation is associated with the accumulation of phosphorylated p53 in mitochondria and the inhibition of mitochondrial metabolism

**DOI:** 10.1101/2023.03.30.534893

**Authors:** Adriana Marques-Carvalho, Beatriz Silva, Francisco B. Pereira, Ha-Neui Kim, Maria Almeida, Vilma A. Sardão

## Abstract

Estrogen deficiency increases bone resorption and is a major contributor to osteoporosis. However, the molecular mechanisms mediating the effects of estrogen on osteoclasts remain unclear. This study aimed at elucidating the early metabolic effects of RANKL – the essential cytokine for osteoclastogenesis – and 17-beta-estradiol (E_2_) on osteoclast progenitor cells, using RAW 264.7 macrophage cell line and bone marrow-derived macrophages as biological models. RANKL stimulated complex I activity, oxidative phosphorylation (OXPHOS), and mitochondria-derived ATP production, as early as 3 to 6 h. This up-regulation of mitochondrial bioenergetics was associated with an increased capacity to oxidize TCA cycle substrates, fatty acids, and amino-acids. E_2_ inhibited all effects of RANKL on mitochondria metabolism. In the presence of RANKL, E_2_ also decreased cell number and stimulated the mitochondrial-mediated apoptotic pathway, detected as early as 3h. Surprisingly, the pro-apoptotic effects of E_2_ were associated with an accumulation of p392S-p53 in mitochondria. These findings elucidate early effects of RANKL on osteoclast progenitor metabolism and suggest novel p53-mediated mechanisms that contribute to postmenopausal osteoporosis.

## Introduction

In the adult skeleton, bone mass is maintained by the continuous turnover of bone in a regulated remodeling process. Derived from hematopoietic progenitor cells, osteoclasts are multinucleated cells highly specialized in resorbing the mineralized bone matrix. Macrophage-colony stimulating factor (M-CSF) and the receptor activator of nuclear factor kappa B ligand (RANKL) are two cytokines critical for the differentiation of osteoclasts from myeloid precursors (Dougall et al., 1999; Teitelbaum & Ross, 2003). RANKL increases the mitochondrial network during osteoclast differentiation and promotes mitochondrial oxygen consumption and ATP production (Arnett & Orriss, 2018; Park-Min, 2019). In the final stages of osteoclast differentiation, RANKL also increases glycolysis and lactate synthesis to meet the high energetic demand of bone resorption (Arnett & Orriss, 2018; Da et al., 2021; Lemma et al., 2016; Taubmann et al., 2020), highlighting the importance of metabolic remodeling during osteoclastogenesis.

Estrogens maintain bone mass and protect the adult skeleton by decreasing osteoclast number via the estrogen receptor (ER) alpha expressed in osteoclastic cells (Almeida et al., 2017; Stavros C. Manolagas, 2000; Hughes et al., 1996). The decrease in estrogen levels after menopause reduces bone mass and strength and increases the risk of fractures (Khosla, 2015; Szulc et al., 2006). One of the mechanisms via which estrogens decrease osteoclast number is promoting their apoptosis (Hughes et al., 1996; Kim et al., 2020). This effect was observed in human and mouse osteoclast precursors cell and RAW 264.7 cells – a mouse monocytic cell line that differentiates into osteoclasts upon stimulation with RANKL (Palacios et al., 2005; Sorensen et al., 2006). The apoptotic effects on osteoclast precursors are significant contributors to the anti-osteoclastogenic effect of estrogens, at least in cell cultures, and are mediated by the pro-apoptotic members of the Bcl-2 family, Bak and Bax (Kim et al., 2020). Following an apoptotic stimulus, cytosolic Bax translocate into the outer mitochondrial membrane, dimerizing with Bak. These Bax/Bak oligomers form pores in the outer mitochondrial membrane, allowing the release of cytochrome C (cyt C) into the cytosol (Hsu et al., 1997), which will promote the assembly of the apoptosome, activation of the caspase pathway, and cell death (Dewson et al., 2012; Zhang et al., 2017). The tumor suppressor p53 is a central regulator of programmed cell death by promoting the transcription of Bax. In addition, stress signals, can phosphorylate p53 at Ser 392 and promote its translocation to mitochondria (Madan et al., 2013). Here, p53 interacts with Bax, promoting its oligomerization and formation of pores, allowing cyt C release and caspase activation (Castrogiovanni et al., 2018; Sardão et al., 2009). P53 can also promote oxidative stress and causes loss of mitochondrial transmembrane electric potential (ΔΨm) due to the opening of mitochondrial permeability transition pore (mPTP) (Bauer & Murphy, 2020). The relevance of p53 activation in the early effects of E_2_ in osteoclast apoptosis is not currently known.

We have previously demonstrated that the effects of RANKL and E_2_ in mitochondria occurred in osteoclast progenitors. RANKL stimulated mitochondrial respiration, while E_2_ prevented this effect after 48 h incubation. The inhibitory effects of E_2_ were associated with a promotion of mitochondria-mediated apoptosis (Kim et al., 2020). In a more recent study, analysis of osteoclastogenesis by single cell transcriptomics revealed an upregulation of genes associated with oxidative phosphorylation, TCA cycle and ETC in pre-osteoclasts after a 24h incubation with RANKL, suggesting that upregulation of mitochondria energetics may be important in the early stages of differentiation (Tsukasaki et al., 2020). Whether this transcriptomic alteration corresponds to early metabolic changes in cells is still unclear. To clarify this issue, we performed a metabolic fingerprinting of the early stages of osteoclast

differentiation. We also examined whether E_2_ impacted the metabolic function during that timeframe and identified p53 as a possible mediator of the effects of E_2_.

## Results

### In the early stages of osteoclast differentiation, E_2_ prevents the metabolic changes induced by RANKL

We have previously shown in bone marrow-derived macrophages (BMMs) that E_2_ abrogates the stimulation of OXPHOS by RANKL at 48 h of exposure (Kim et al., 2020). Here, we first investigated the timeline of E_2_ effects on OXPHOS by measuring the cellular oxygen consumption ratio (OCR), using the Seahorse Xf^e^96 Extracellular Flux Analyzer. To this end, OCR in RAW 264.7 cells was assessed after stimulation with RANKL for three distinct early time points, 3, 6, and 24 h, in the presence or absence of E_2_ (Figure 1a). RANKL stimulated basal, maximal, and ATP production-linked OCR (Figure 1b-d). RANKL also increased proton leak and non-mitochondrial oxygen consumption at 6 and 24 h time points, respectively (Supplementary Figure S1a-b). We next examined whether RANKL altered glycolysis by measuring the extracellular acidification rate (ECAR), but no significant alterations in ECAR were observed at any tested time point (Supplementary Figure S1c). In line with the distinct effects of RANKL on OXPHOS and glycolysis, the rate of mitochondrial ATP production is highly increased at 6 and 24 h while the rate of glycolytic ATP production is only modestly increased at 24 h (Figure 1e-f). Further, an increase in total ATP levels (Figure 1g) at all tested time points was also observed. The presence of E_2_ prevented the increase in cellular OCR, total ATP levels, and the rate of mitochondrial ATP production induced by RANKL at all tested time points.

**Figure 1.**
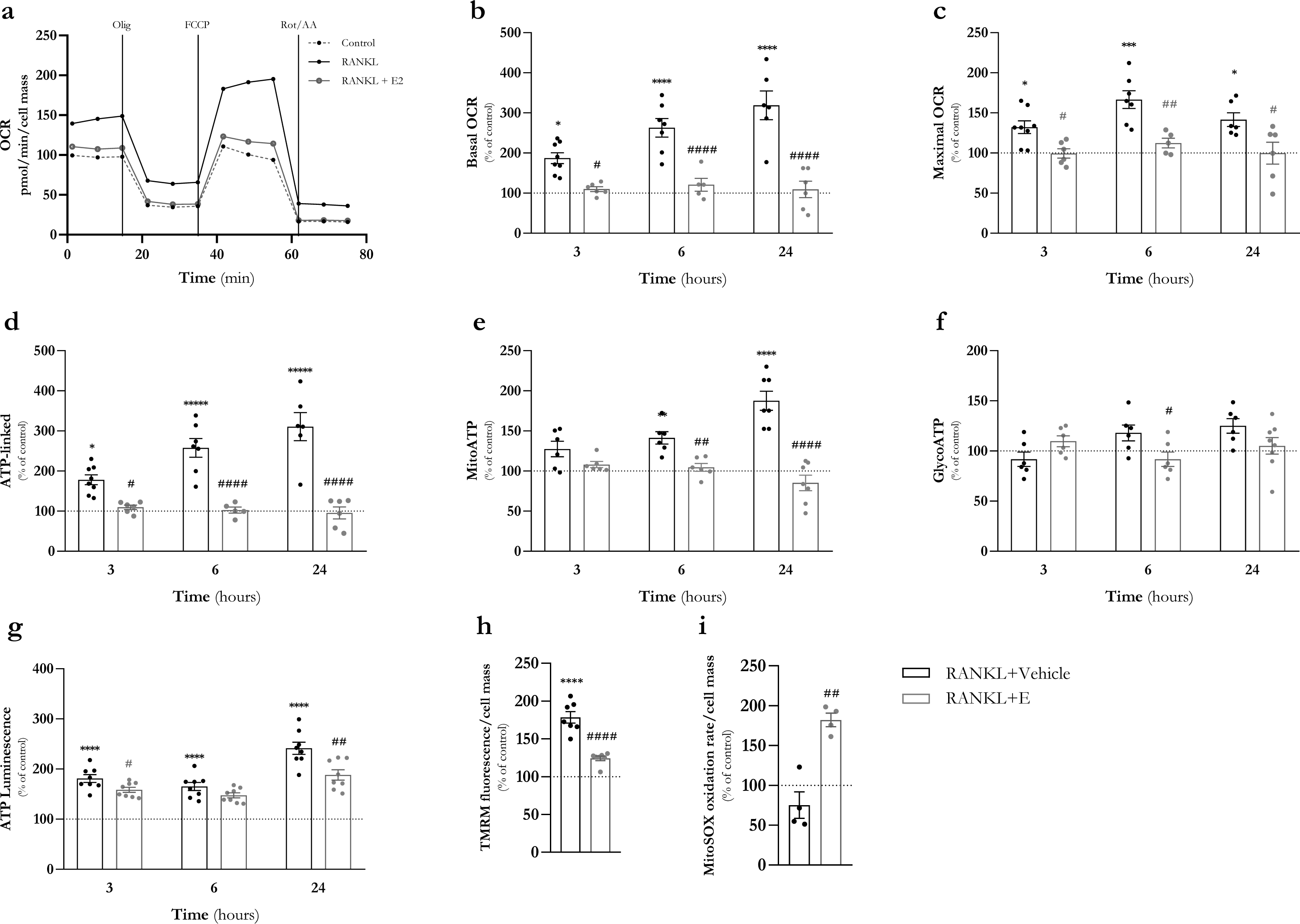
E_2_ decreases oxygen comsumption rate, ATP production, mitochondrial membrane potential and ROS production at the early stages of osteoclast differentiation in RAW 264.7 cells. RAW 264.7 cells were treated RANKL (30 ng/mL) with or without E_2_ (10^-8^ M) for 3, 6 and 24 h. Typical representation of oxygen consumption rate (OCR) measurements **(A)**. Several respiratory parameteres were evaluated: **(B)** basal OCR, **(C)** maximal OCR and **(D)** ATP-linked OCR. **(E-F)** RAW 264.7 cells treated with RANKL and E_2_ for 24 h and the rate of glycolitic and mitochondrial ATP production was quantified using the Real-time ATP Rate Seahorse assay. **(G)** ATP levels were determined by CellTiter-Glo Luminescent Cell Viability Assay after 3, 6 and 24 h incubation with RANKL and E_2_. **(H)** Mitochondrial membrane potential and **(I)** Mitochondrial ROS alterations after 3 h treatment with RANKL in the presence or absence of E_2_. Data are mean ± SEM of 4-8 biological replicates. Each dot represents one replicate. The results were normalized for the control condition (cells without treatment, 100% marked by a doted line). Control vs RANKL (black) and RANKL vs RANKL+E_2_ (grey) significance was accepted with p<0.05, using two-way ANOVA analysis with Tukey’s multiple comparisons test.

Following, we examined whether the same early effects of RANKL and E_2_ would also be observed in BMMs cells. As observed for RAW 264. 7 cells, RANKL stimulated cellular OCR (except proton leak) and the rate of mitochondrial ATP production in BMM cells after 6 and 24 h and the presence of E_2_ attenuated all these effects (Figure 2a-g and Supplementary Figure S2a-c). E_2_ alone had no effect on any of the measured parameters in both RAW 264.7 cells and BMMs (Supplementary Figure S1d and Figure S2d).

**Figure 2.**
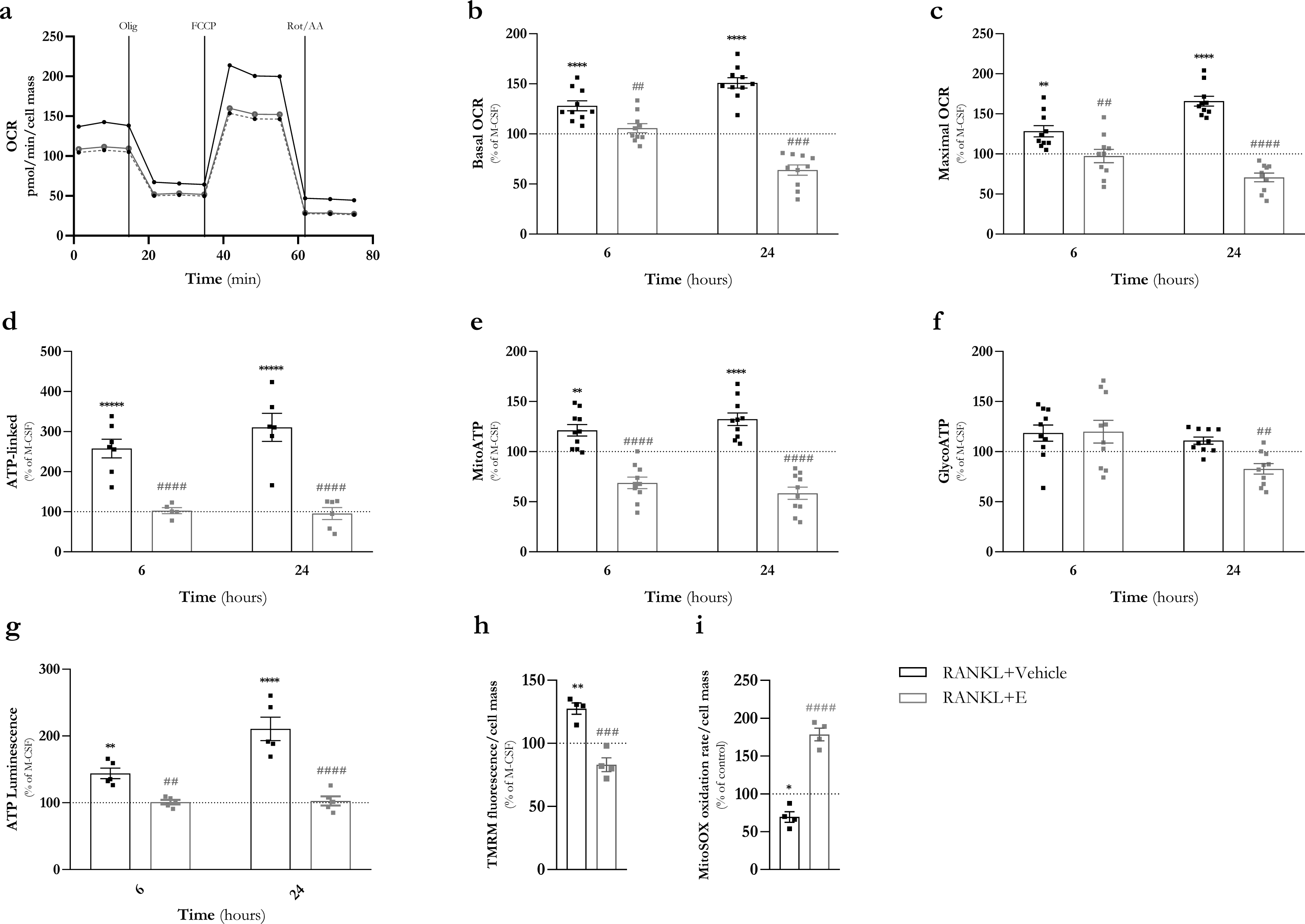
E_2_ induces mitochondrial dysfunction at the early stages of osteoclast differentiation in BMMs. BMMs cells were suplemented with M-CSF (30 ng/mL) and treated RANKL (30 ng/mL) with or without E_2_ (10^-8^ M) for 6 and 24 h. Typical representation of oxygen consumption rate (OCR) measurements **(A)**. Several respiratory parameteres were evaluated: **(B)** basal OCR, **(C)** maximal OCR and **(D)** ATP-linked OCR. **(E-F)** RAW 264.7 cells treated with RANKL and E_2_ for 24 h and the rate of glycolitic and mitochondrial ATP productin was quantified using the Real-time ATP Rate Seahorse assay. **(G)** ATP levels were determined by CellTiter-Glo Luminescent Cell Viability Assay after 3, 6 and 24 h incubation with RANKL and E_2_. **(H)** Mitochondrial membrane potential and **(I)** Mitochondrial ROS alterations after 3 h treatment with RANKL in the presence or absence of E_2_. Data are mean ± SEM of 3-10 independent experiments. Each dot represents one animal. The results were normalized for the control condition (cells with M-CSF, 100% marked by a doted line). M-CSF vs RANKL (black) and RANKL vs RANKL+E_2_ (grey) significance was accepted with p<0.05, using two-way ANOVA analysis with Tukey’s multiple comparisons test.

A decrease in ΔΨm and increased production of mitochondrial reactive oxygen species (ROS) are characteristics associated with mitochondrial dysfunction (Kuznetsov et al., 2011). To assess whether the inhibitory effects of E_2_ on mitochondrial respiration were associated with a decrease in ΔΨm and increased ROS production, we evaluated ΔΨm and mitochondrial superoxide anion in both RAW 264.7 cells and BMMs after 3 or 6 h of exposure to RANKL and RANKL plus E_2_. Compared with untreated cells, RANKL increased ΔΨm in both models but did not affect mitochondrial superoxide anion. E_2_ attenuated ΔΨm induced by RANKL and increased mitochondrial superoxide anion levels, suggesting that E_2_ causes mitochondrial alterations (Figure 1h-i and Figure 2h-i).

We then applied a decision tree classifier to samples from RAW 264.7 cells treated with RANKL and RANKL plus E_2_ at 3, 6, and 24 h timepoints. The computational analysis aims to gain insight into the effects of RANKL and estrogens on mitochondria and to assess if it is possible to separate the samples considering just the information provided by mitochondrial respiration and ATP production subsets. The effect of E_2_, in all three time points, was robust enough for a perfect separation between conditions (Supplementary Figure S3a). The confusion matrix obtained with test data confirmed this separation (Supplementary Figure S3b). The study was repeated with BMM, for 6 and 24 h timepoints. The resulting model has 4 decision levels (Supplementary Figure S3c), suggesting a slightly increased difficulty in defining the boundaries that separate the two studied conditions. Once again, maximal respiration is the most relevant feature for classification.The test accuracy was around 95%, as confirmed by the data displayed in the confusion matrix (Supplementary Figure S3d). The analysis of the results unveils the existence of a high discriminative inhibitory effect of estrogens on osteoclasts, which correlates with the biological outcome.

### E_2_ decreases the levels of OXPHOS subunits, complex I activity and cardiolipin levels

To determine if the changes observed in mitochondrial respiration and ATP production were associated with altered content of mitochondrial respiratory chain complexes, we evaluated protein levels of subunits of OXPHOS complex I (NDUFB8), complex III (UQCRC2), complex IV (MTCO1), and complex V (ATP5A) (Figure 3a-b). After 6 h of treatment, RANKL increased the levels of NDUFB8 subunits, compared to untreated cells but did not alter the evaluated subunits from the other complexes (Figure 3a-b). E_2_ prevented the effect of RANKL on NDUFB8 subunits and decreased ATP5A subunit.

**Figure 3.**
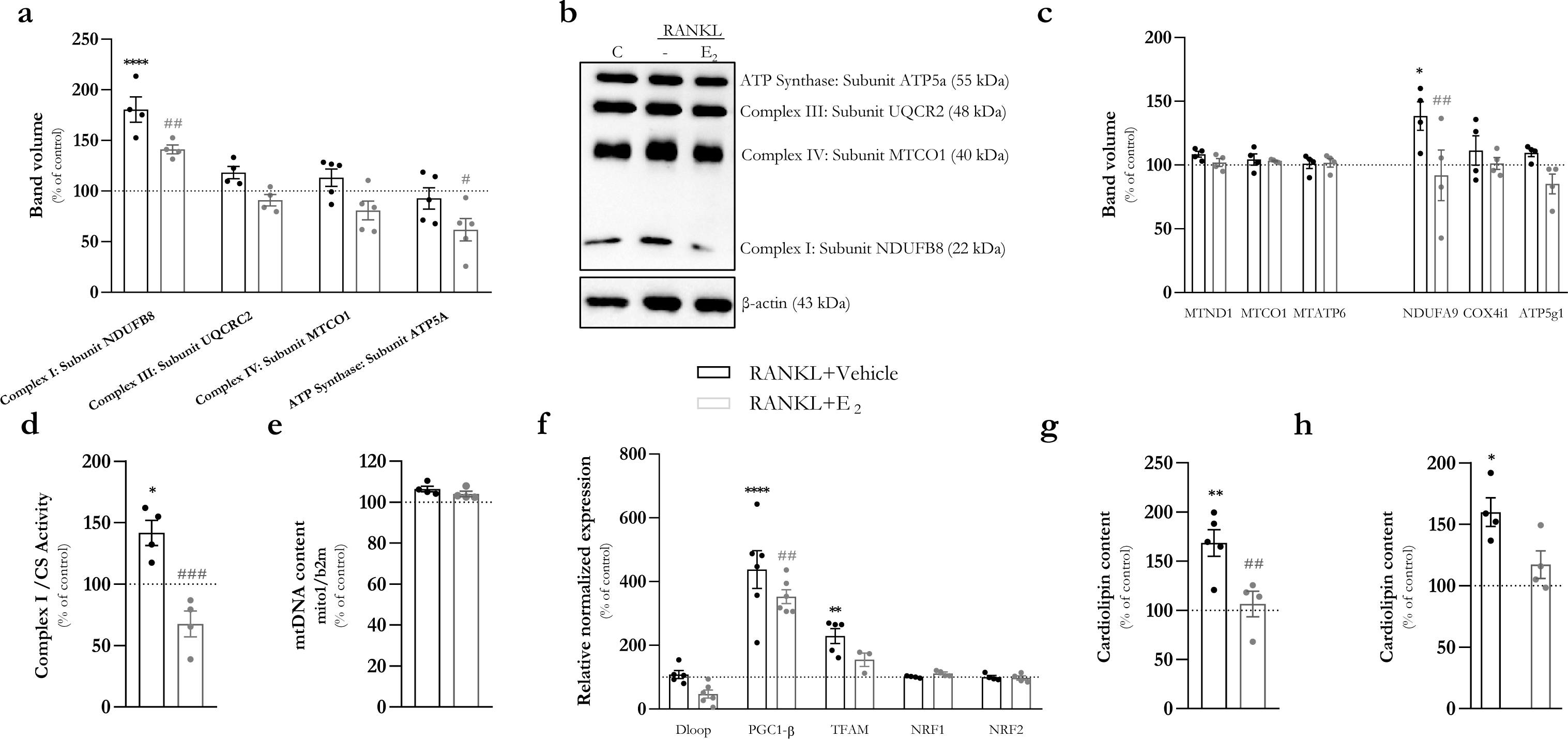
Estrogen decreases OXPHOS protein levels, complex I activity and mitochondrial biogenesis in RAW 264.7 cells. RAW 264.7 cells, after 3 h incubation with RANKL (30 ng/mL) with or without E_2_ (10^-8^ M)**. (A-B)** Western blot of mitochondrial OXPHOS protein levels: NDUFB8 (complex I), SDHB (complex II), UQCRC2 (complex III), COXII (complex IV), ATP5A (complex V) subunits. The levels were normalized to β-actin levels and for the control group (cells without treatment, 100% marked by a doted line). **(C)** Changes on the transcripts of nuclear and mitochondrial-encoded oxidative phosphorylation complexes proteins. **(D)** Complex I (NADH dehydrogenase) activity was determined according to a previously described protocol using 10 μg of protein (Spinazzi et al., 2012). Citrate synthase activity was used to normalize complex I activity and the final results were normalized to the control condition (cells without treatment, 100% marked by a doted line). **(E)** mtDNA copy number was evaluated by qPCR, based on the amplification of mito1 and beta-2-microglobulin (b2m); **(F)** Changes on the transcripts of proteins associated with mitochondrial biogenesis (DLoop, PGC1-β, Tfam, NRF1 and NRF2) evaluated by qRT-PCR; **(G)** Cardiolipin content evaluated by Cardiolipin Assay kit (MAK362, Sigma, Merck) and **(H)** Cardiolipin Synthase 1 (CRLS1) levels. Data are expressed as mean ± SEM of 3-6 independent experiments. Control vs RANKL (black) and RANKL vs RANKL+E_2_ (grey) significance was accepted with p<0.05, using two-way ANOVA analysis with Tukey’s multiple comparisons test.

We have previously shown that BMM lacking ERα exhibit increased expression of several nuclear genes encoding mitochondrial complex I subunits. Moreover, complex I activity is increased after 6 h of exposure to RANKL, with this effect attenuated by E_2_ (Kim et al., 2020). Similarly to BMMs, RANKL also stimulated the expression of nuclear-encoded complex I NDUFA9 subunit, but did not affect the expression of nuclear-encoded subunits of other complexes or the mitochondrial-encoded subunits (Figure 3c). Moreover, RANKL increased complex I activity in RAW 264.7 cells after 3 h, and these effects were prevented by E_2_ (Figure 3d). In contrast, RANKL and E_2_ did not affect mitochondrial DNA (mtDNA) copy number at this early time point (Figure 3e), suggesting that these early effects on OXPHOS were not due to changes in mtDNA. Consistent with these findings, the D-loop content, a mitochondrial region responsible for the regulation of mtDNA replication and transcription (Takamatsu et al., 2002), was not altered (Figure 3f).

We also examined the mRNA expression of factors responsible for mitochondria biogenesis, such as Nrf1, Nrf2, TFAM, and PGC1β (Gureev et al., 2019; Shao et al., 2010). We found no changes in the amount of Nrf1 and Nrf2, in RAW 264.7 cells after 3 h incubation with RANKL, in the presence or absence of E_2_. RANKL increased the expression of PGC1β and TFAM, while E_2_ attenuated the effect of RANKL in PGC1β expression (Figure 3f).

Cardiolipin (CL) is a specific mitochondrial phospholipid that plays a central role in the assembly of respiratory supercomplexes and mitochondrial biogenesis (Dudek, 2017). We evaluated total CL and cardiolipin-synthase (CRLS) 1 in response to RANKL and E_2_ treatments. RANKL increased the amount of both CL and CRLS after 3 h, whereas E_2_ inhibited this effect (Figure 3 g-h).

### E_2_ decreases mitochondrial metabolic substrate consumption

To further characterize the effects of RANKL and E_2_ on mitochondrial metabolism, we measured the oxidation of 23 different metabolic substrates during the early stages of osteoclastogenesis. The assay was performed using permeabilized RAW 264.7 cells and BMMs previously cultured with RANKL and E_2_ for 24 h. In both models, cells treated with RANKL were able to oxidize substrates from the TCA cycle, fatty acids, amino acids, and other substrates such as γ-amino-butyric acid and α-keto-isocaproic acid (Figure 4a). The addition of E_2_ reduced the capacity of both cellular models to oxidize the substrates from all the mentioned energy pathways (Figure 4a). A visual comparison between the capacity to oxidize TCA cycle substrates (Figure 4b), fatty acids (Figure 4c), and amino acids (Figure 4d) is also provided by a set of radar charts. For all different pathways and individual substrates, the average values in RANKL-treated cells are always higher than the untreated cells. In the presence of E_2_ this trend was reverted.

**Figure 4.**
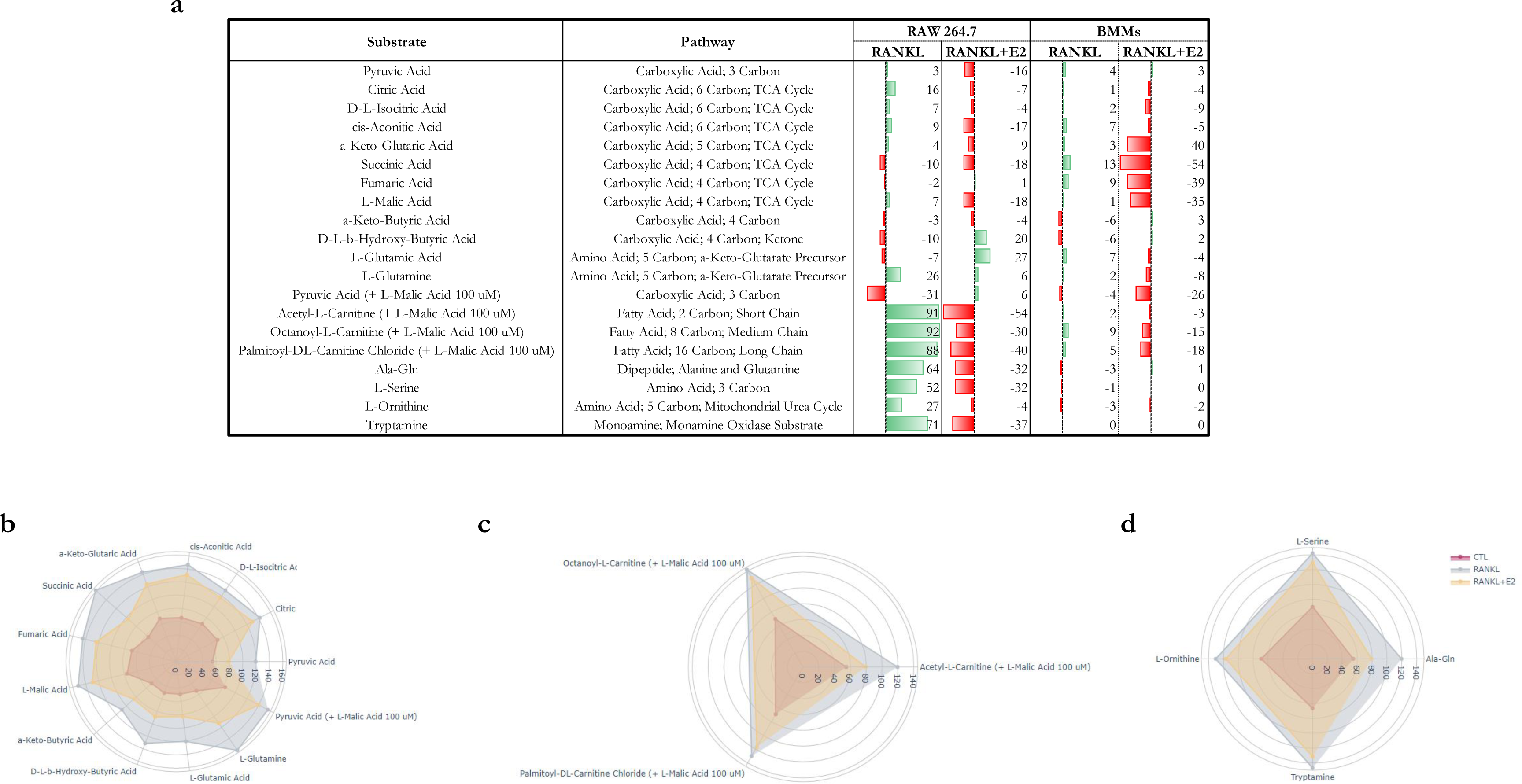
E_2_ decreases mitochondrial metabolism. RAW 264.7 cells and BMMs were treated with RANKL (30 ng/mL) with or without E_2_ (10^-8^ M) for 24 h. **(A)** The rate of oxidation of different substrates from different pathways (TCA cycle, fatty acids and aminoacids) was determined using Biolog Mitoplase S1 assay. Radar charts representing the multivariate date of RAW 264.7 cells, as an average value for **(B)** TCA cycle, **(C)** fatty acids and **(D)** aminoacids substrates. Data is represented as a mean of 3 independent experiments for RAW 264.7 or 1 independent experiments for BMMs..

### E_2_ induces early apoptotic events

We have previously shown in BMM that incubation with E_2_ during the first 24 h of culture was sufficient to decrease the number of osteoclasts after a 5-day culture (Kim et al., 2020). This decrease is associated with an apoptotic effect of estrogen via the mitochondrial death pathway. Here, we examined whether RANKL and E_2_ had effects on RAW 264.7 cell number after 3, 6, 12, and 24 h. RANKL increased cell number after 24 h treatment (Figure 5a). E_2_ alone had no effect on cell number, but in the presence of RANKL decreased cell number after 3 h treatment. To determine whether the effects of E_2_ on cell number were associated with pro-apoptotic events, we initially analyzed Bax and Bak protein levels in mitochondria-enriched fractions after 3 h treatment. RANKL alone had no effects on the levels of mitochondrial Bax and Bak but in the presence of E_2_ an increase in the levels of both proteins (Figure 5b-c) was observed at this early time point. In line with these changes, the presence of E_2_ stimulated cyt C release from mitochondria to the cytosol (Figure 5d-e) and increased Caspase 9 activity after 6 h of treatment (Figure 5f). These changes were followed by an increase in the effector Caspase-3 activity after 12 h. (Figure 5g). On the other hand, treatment with RANKL or E_2_ alone had no impact on caspase activity (Figure 5h).

**Figure 5.**
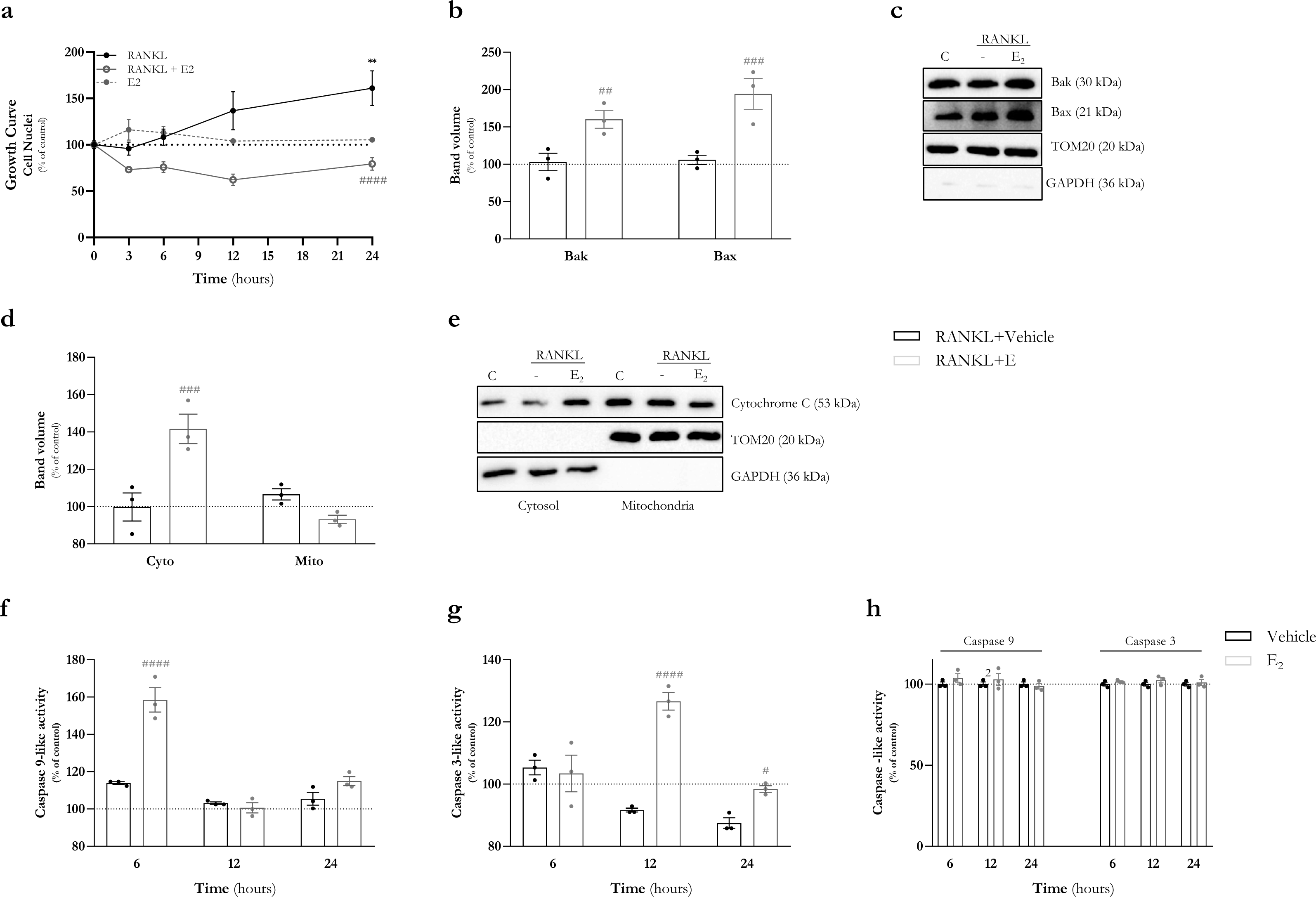
E_2_ decreases cell number and induce rapid cell apoptotic events. RAW 264.7 cell growth evaluated by **(A)** nuclei counting by Hoechst 33342 staining after incubation with RANKL (30 ng/mL), E_2_ (10^-8^ M) or RANKL plus E_2_ for 3, 6, 12 or 24 h. **(B-C)** Western blot of Bax and Bak protein levels in mitochondrial-enriched fraction of RAW 264.7 cells, after 3 h incubation with RANKL (30 ng/mL) with or without E_2_ (10^-8^ M). The levels were normalized to TOM20 levels in the mitochondrial and for the control group (cells without treatment, 100% marked by a doted line). **(D-E)** Western blot of cytochrome C protein levels in cytosolic and mitochondrial-enriched fractions, after 6 h incubation with RANKL with or without E_2_. The levels were normalized to TOM20 levels in the mitochondrial fraction or GAPDH levels for the cytosolic fraction and for the control group (cells without treatment, 100% marked by a doted line). **(F)** Caspase 9-like and **(G)** Caspase 3-like activity in RAW 264.7 cells after 6, 12 h and 24 h incubation, with RANKL (30 ng/mL) and E_2_ (10^-8^ M). **(H)** Caspase 9-like and caspase 3-like activities of RAW 264.7 cells were treated with or without E_2_ (10^-8^ M) for 3, 6 and 24 h. Data is represented as a mean of 3 independent experiments for RAW 264.7. Control vs RANKL (black) and RANKL vs RANKL+E_2_ (grey) significance was accepted with p<0.05, using two-way ANOVA analysis with Tukey’s multiple comparisons test.

### The presence of E2 during osteoclastogenesis promotes P53 phosphorylation at Ser 392 and P53 translocation to mitochondria

Our previous finding showed that E_2_ stimulated all the different steps of the mitochondria pathway of apoptosis, including an increase in Bax in the mitochondria, release of cytochrome c into the cytosol and activation of caspases. The tumor suppressor p53 is a key factor responsible for cell fate determination by controlling cell-cycle arrest, apoptosis, and cellular senescence. p53 is a short-lived protein whose the levels of which are tightly controlled by Mdm2 E3 ligase (Brooks and Gu, 2006; Haupt et al., 1997; Kubbutat et al., 1997). The majority of p53 resides in the nucleus, where it can bind to the promoter and/or enhancer of its target genes. Upregulation of Puma, Bax, and Noxa is thought to mediate p53-dependent apoptosis (Miyashita et al., 1994; Nakano and Vousden, 2001; Oda et al., 2000). In response to stress stimuli, p53 also moves to the mitochondrial outer membrane where it binds to BAK/Bax to promote BAK/Bax oligomerization (Chipuk et al., 2004; Mihara et al., 2003).

In some cell types, p53 gene is under the control of ER-α transcriptional activity (Berger et al., 2012). Because of this evidence, we examined whether p53 was involved in the pro-apoptotic effects of E_2_ in osteoclast precursors. No changes in the total levels of p53 were observed after 3 h of treatment (Figure 6a-b). In contrast, the levels of p53 phosphorylation at Ser 392 were increased by E_2_ in the mitochondria-enriched fractions (Figure 6a-b). The magnitude of the increase of p53 phosphorylation was higher in the mitochondrial fractions.

**Figure 6.**
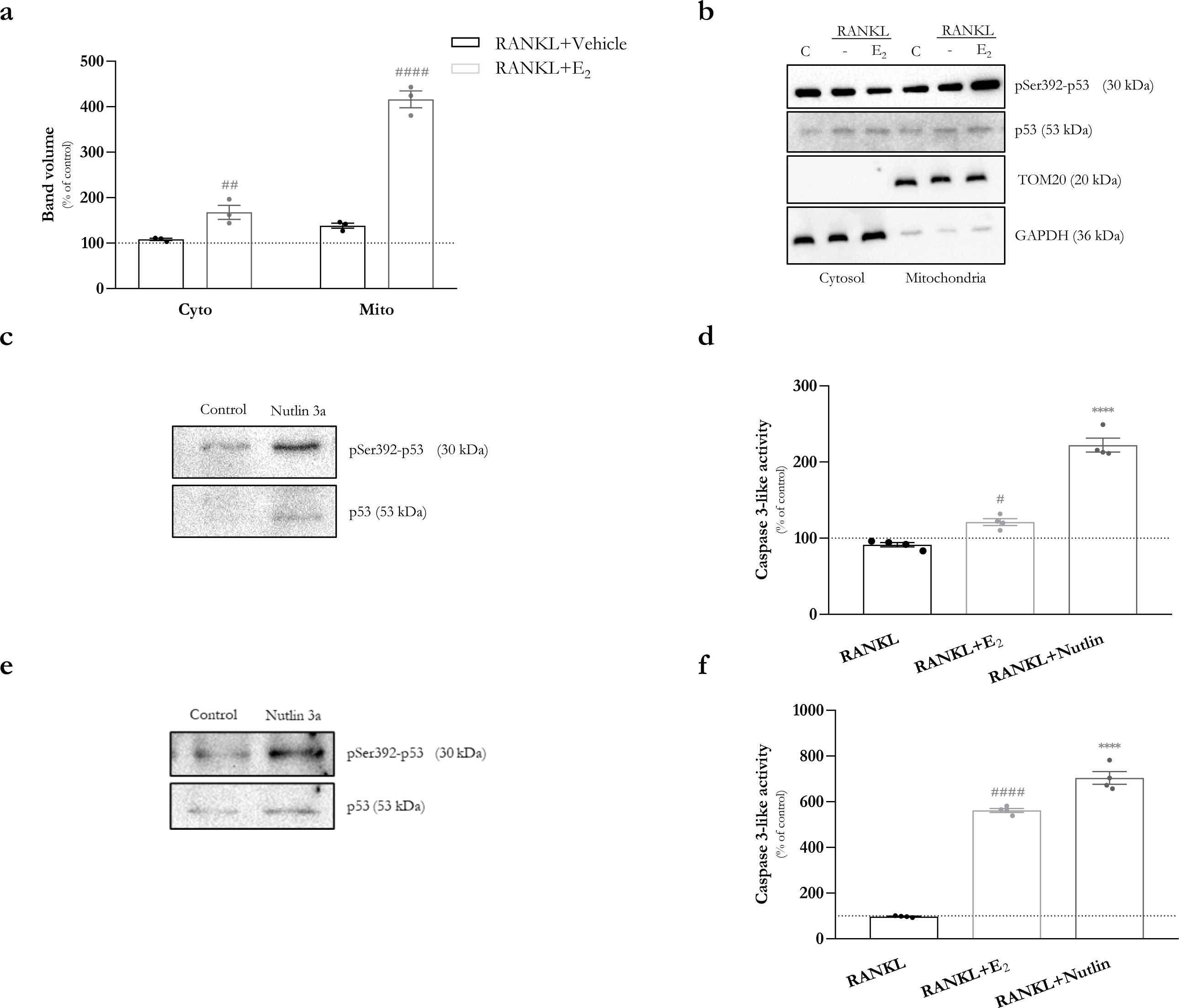
Estrogen induces P53 phosphorylation and translocation to mitochondria. **(A-B)** Western blot of phosphorylated p53 (Ser 392) and p53 protein levels in cytosolic and mitochondrial-enriched fractions, after 3 h incubation with RANKL with or without E_2_. The levels were normalized to TOM20 levels in the mitochondrial fraction or GAPDH levels for the cytosolic fraction and for the control group (cells without treatment, 100% marked by a doted line).Western blot of phosphorylated p53 (Ser 392) and p53 in total cellular extracts and caspase 3-like acitivty of RAW 264.7 **(C-D)** and BMMs **(E-F)** incubated with alone with E_2_ (10^-8^ M), RANKL (30 ng/mL) plus E_2_ or nutlin-3a (10 μM) for 12 or 24 h, respectively. The levels were normalized to β-actin levels and for the control group (cells without treatment, 100% marked by a doted line). Data are expressed as mean ± SEM of 3-4 independent experiments. Control vs RANKL (black) and RANKL vs RANKL+E2 (grey) significance was accepted with p<0.05, using one-way or two-way ANOVA analysis with Tukey’s multiple comparisons test. In Nutlin-3a experiments, RANKL+Nutlin vs RANKL was accepted with p<0.0001.

We next examined whether activation of p53 independently of E_2_ could mimic the apoptotic effects of E_2_. To this end, we incubated RAW 264.7 and BMMs with 1 µM of nutlin-3a for 12 and 24 hours, matching the maximal caspase 3 activity induced by E_2_ determined in each cell type [Figure 4 and (Kim et al., 2020)].

Nutlin-3a is a small molecule that inhibits p53 ubiquitination by inhibiting the interaction between MDM2 and p53, consequently inducing cell autophagy and apoptosis. Our results showed that Nutlin-3a increased total p53 and pSer392-p53 protein levels in RAW 264.7 (Figure 6c) and BMMs (Figure 6d), promoting caspase-3 activation in both models (Figure 6e-f).

## Discussion

The differentiation of macrophages into bone-resorbing osteoclasts is accompanied by changes in energy metabolism and an increase in mitochondria mass (Arnett & Orriss, 2018; Indo et al., 2013; Ishii et al., 2009; Park-Min, 2019). This increase in mitochondria mass has been described after 48h exposure to RANKL, coinciding with the initiation of cell fusion (Ishii et al., 2009; Zeng et al., 2015). We have recently uncovered a stimulatory effect of RANKL on mitochondria of osteoclast progenitors, which is inhibited by estrogens. These changes precede the increase in mitochondrial mass. We have also proposed that the effects of E_2_ on mitochondria might be responsible for the pro-apoptotic action of estrogens at the early stages of osteoclast differentiation (Kim et al., 2020). There is no data on the early responses of osteoclast mitochondria to E_2_ and how that shapes cell fate. Here, for the first time we examined in detail the early actions of RANKL and E_2_ on osteoclast metabolism and cell death.

Our findings in the present paper show that RANKL stimulated OXPHOS (basal, maximal, and ATP-linked respiration) and mitochondrial ATP production as early as 3 to 6 h, in RAW 264.7 cells and primary bone marrow-derived macrophages. As seen before, E_2_ inhibited these rapid effects of RANKL at later time points. The effects of RANKL and E_2_ on mitochondrial metabolism were confirmed by computational analysis, in which a decision tree was trained by using OCR seahorse data, from both RAW 264.7 cells and BMMs. Thus, these effects on mitochondria are initiated much earlier than 48 h, as we previously described (Kim et al., 2020). Also, following our previous study (Kim et al., 2020), E_2_ had no effect on mitochondria OXPHOS in macrophages in the absence of RANKL. Thus, the effects of E_2_ are likely to result from the interference with a signal initiated by RANKL to stimulate mitochondria activity and not from direct actions of E_2_ on mitochondria. Currently, the signals responsible for the early effects of RANKL on mitochondria remain unknown.

Using a genetic unbiased approach, we found that ERα signaling in osteoclast progenitors attenuates the expression of several proteins of mitochondrial complex I, and that E_2_ suppresses the activity of complex I promoted by RANKL (Kim et al., 2020). Studies in a mouse model in which complex I in macrophages is deficient indicate that complex I is required for normal bone resorption. In line with the rapid effects on OXPHOS and ATP production, we show here in RAW 264.7 cells that the stimulatory actions of RANKL on complex I and inhibition by E_2_ occurs as early as after 3 h of exposure. Together, these findings support the notion that the early stimulation of complex I activity by RANKL promotes osteoclastogenesis, and inhibition of these effects contributes to the anti-osteoclastogenic actions of estrogens.

The early up-regulation of mitochondrial bioenergetics by RANKL was associated with an increased capacity to oxidize TCA cycle substrates, fatty acids, and amino-acids. Similar to the effects on OXPHOS, E_2_ prevented energy substrate consumption by RANKL. In agreement with our findings, proteomic studies have suggested that fatty acid and amino acid oxidation are up-regulated in mature osteoclasts (An et al., 2014). In contrast to the stimulatory effects on mitochondria, RANKL had minimal, if any, effect on glycolysis and glycolytic ATP production at the early time points in both RAW 264.7 cells and BMMs. Together, these results support the idea that during the early stages of osteoclastogenesis cells rely on mitochondrial energy-producing pathways. Indeed, others have previously suggested that OXPHOS is the prevalent energy pathway during early stages of osteoclastogenesis, while glycolysis is up-regulated at a later stage, in mature osteoclasts (Indo et al., 2013) (Taubmann et al., 2020).

In this report, we show that in RAW 264.7 cells, neither RANKL nor E_2_ affected mtDNA content, suggesting that the metabolic effects we observed were independent of changes in mitochondrial mass. Nonetheless, RANKL increased the expression of genes associated with mitochondrial biogenesis, such as Tfam and PGC1-β while NRF1 and NRF2 were not affected. Moreover, RANKL increased CRLS1 levels and cardiolipin content. These actions, most likely, are aimed at preparing a subsequent expansion of the mitochondria network in response to RANKL, as previously described in the primary osteoclast (Lemma et al., 2016). E_2_ prevented all the stimulatory effects of RANKL on mitochondrial genes and cardiolipin.

We have previously shown that E_2_ induces apoptosis of bone marrow-derived osteoclast progenitors and thereby decreases osteoclast formation via a Bak and Bax-dependent mechanism (Kim et al., 2020). We found here that this mitochondria-dependent apoptotic pathway is stimulated by E_2_ as early as 3 h, as indicated by the accumulation of Bax in the mitochondria and subsequent release of cyt C to the cytosol and activation of caspase-9 and caspase-3.

The pro-apoptotic effect of estrogen was also associated with the accumulation of phospho-p53 in mitochondria. P53 phosphorylation at Ser-392 stabilizes p53 and induces its mitochondrial translocation after a stress stimulus. In mitochondria, p53 can promote apoptosis by interacting and stimulating Bax oligomerization, and inducing cyt C release (Castrogiovanni et al., 2018). Our findings show that the accumulation of p53 in mitochondria coincides with cyt C release, supporting the idea that this effect contributes to the pro-apoptotic actions of E_2_ in osteoclast progenitors. In line with these results, stimulation of p53 with Nutlin 3a, a potent and selective Mdm2 antagonist that promotes p53 activity, mimicked the effects of estrogen.

P53 functions largely as a transcription factor and can be activated in response to a variety of cellular stresses (Kruiswijk et al., 2015). Once activated can regulate an array of pathways, including cell cycle arrest, programmed cell death, DNA repair or metabolism (Green & Kroemer, 2009; Riley et al., 2008). The action of p53 differs between cell type and is not evident how to predict the propensity of the different cell types to undergo apoptosis versus other responses, in response to p53 activation (Kracikova et al., 2013; Rizzotto et al., 2020). In osteoclasts precursors, Nutlin promoted the activation of programmed cell death, suggesting that, in osteoclast precursors, p53 stimulates apoptosis.

In addition, E_2_ might also stimulate p53 nuclear activity as indicated by the increase in Bax levels. Mice lacking p53 have an increased number of osteoclast and bone resorption (Wang et al. JCB 2006). However, osteoclastogenesis is unaffected in cultured bone marrow cells from these mice. It was suggested that the increased osteoclasts in p53^-/-^ mice resulted from increased production of M-CSF by osteoblastic cells. In view of our findings here, a plausible explanation for the increase in the number of osteoclasts in p53^-/-^ mice could be an attenuation of the anti-osteoclastogenic effects of estrogen. Interestingly, in breast cancer cells, E_2_ via ERα represses p53 mediated transcription (Berger et al., 2012), highlighting yet again the distinct effects of E_2_ in osteoclasts.

In summary, we show that that anti-osteoclastogenic effects of E_2_ in osteoclast progenitors are initiated very early in the differentiation process and are similar in both RAW 264.7 cell line and primary bone marrow derived macrophages. Interestingly, we found for the first time that the early effects of E_2_ are associated with accumulation of ser392-phosphorylated p53 in the mitochondria, as well as with an inhibition of cellular metabolism. Our observations indicate that the mitochondria apoptotic pathway, mediated by Bax and Bak, seems to be stimulated by p53, and plays a critical role in the anti-osteoclastogenesis actions of estrogens. The significance of this pathway in osteoclasts is highlighted by the evidence that mice lacking Bax and Bak present multiple anomalies in bone marrow myeloid cells number. The work presented here underlies the contribution of mitochondria in osteoclast physiology and pathophysiology and suggests a new signal that leads to Bak and Bax activation in response to estrogens, which could be a new pathophysiological mechanism for the development of postmenopausal-associated osteoporosis.

## Materials and Methods

### Cell culture and cell treatments

*Mus musculus* RAW 264.7 cell line (TIB-71, Lot n. ° 62654190, ATCC®, Manassas, VA) was cultured with Dulbecco’s Modified Eagle’s Medium (DMEM; D5030, Sigma-Aldrich, St. Louis, MO, USA) supplemented with 25 mM D-(+)-Glucose, 4 mM L-Glutamine, 0.1 mM NaH_2_PO_4_.H_2_O, 18 mM Sodium Bicarbonate, 1 mM Sodium Pyruvate, 10% fetal bovine serum (FBS) and 1% penicillin/streptomycin in a humidified atmosphere (5% CO_2_, 37°C). Cells were used from the 8^th^ until the 18^th^ passage to avoid loss of ability to differentiate (Taciak et al., 2018). To evaluate the early effects of estrogen in osteoclast differentiation, cells were seeded at 6x10^4^ cells/cm^2^ and grown for 24 h until they reached 70-80% confluency and following treated with RANKL (30 ng/mL; 462-TR, R&D systems), with or without E_2_ (10^-8^ M; E1024, Sigma-Aldrich, St. Louis, MO, USA) for 3, 6, 12 or 24 h, depending on the assays.

Bone marrow-derived macrophages (BMMs) were obtained, as previously described, from 10 old male C57BL/6J mice at 4-6 months of age, obtained from Charles River Laboratories France S.A.S. (Charles River, Barcelona, Spain). Animals were housed under controlled 12 h light/dark cycles at 20-24°C with 45-65% of humidity. Whole bone marrow cells were flushed from femurs and tibias, and ACK buffer was used to deplete red blood cells. Cells were plated in α-MEM (11900-024, Thermo Fisher Scientific, Hampton, NH, USA), supplemented with 10% FBS and 1% penicillin/streptomycin with macrophage-colony stimulating factor (M-CSF, 30 ng/mL; 416-ML, R&D systems, Minneapolis, MN, USA) for 4 days to obtain BMMs, which were used as osteoclast precursors. Cells were seeded at of 15x10^4^ cells/cm^2^, grown for 24 h and treated with RANKL (30 ng/mL; 462-TR, R&D systems, Minneapolis, MN, USA), with or without E_2_ (10^-8^ M; E1024, Sigma-Aldrich, St. Louis, MO, USA) for 3, 6 or 24 h, depending on the assays. The use of animal models was approved by the Animal Welfare Committee at the University of Coimbra (ORBEA_131_2016/24032016) and by the Portuguese Authority of Directorate-General for Food and Veterinary (DGAV - 0421/000/000/2016). All the procedures were also conducted in accordance with the European Union directive (2010/63/EU) by accredited users.

### Cellular oxygen consumption and extracellular acidification rate measurements

The cellular oxygen consumption and extracellular acidification rates (OCR and ECAR, respectively) were measured using the Seahorse XFe96 Extracellular Flux Analyzer (Agilent Technologies, Santa Clara, CA, USA). Seahorse XF Cell Mitostress test and Real-Time ATP Rate Assays (Agilent Technologies, Santa Clara, CA, USA) were used to evaluate mitochondrial function. Briefly, RAW 264.7 cells and BMMs were plated in a Seahorse XF Cell Culture Microplate and treated with RANKL (30 ng/mL) with or without E_2_ (10^−8^ M) for 3, 6 and 24 h. The day before the assay, the XFe96 sensor cartridge was hydrated with 200 µl of calibration buffer and left overnight at 37°C. On the day of the assay, low-buffered serum-free minimal DMEM (assay medium) was prepared by supplementing DMEM D5030 with 25 mM D-(+)-Glucose, 4 mM L-Glutamine, 1 mM Sodium Pyruvate and 5 mM of HEPES, pH 7.4. After treatment, the cell culture medium was replaced by 175 μL of pre-warmed assay medium, and the cells were incubated at 37°C for 1 h in a non-CO_2_ incubator. For Mitostress Assay, oligomycin (3 µM for RAW 264.7 and BMMs), FCCP (0.25 µM for RAW 264.7 and 2 µM for BMMs) and rotenone plus antimycin A (2 µM for RAW 264.7 and BMMs) were sequentially injected through the ports A, B and C of the XFe96 sensor cartridge. For the Real-Time ATP Rate Assay of RAW 264.7 cells, 3 µM oligomycin were injected through Port A, followed by the injection of 2 µM rotenone plus 2 µM antimycin A through port B. Three baseline rate measurements using a 3 min mix and 3 min measure for 3 cycles were used to measure baseline OCR and ECAR levels, followed by 3 min mix and 3 min measure for 3 cycles after each injection. The assays were normalized by SRB staining signal (section 2.7) and analyzed using the Software Version Wave Desktop 2.6 (Agilent Technologies, Santa Clara, CA, USA). At the end of the assay, the cells were fixed overnight with 10% TCA at 4°C for further SRB normalization (section 2.6).

### Intracellular ATP levels determination

Intracellular ATP levels were measured by a lucifrin-lucifrase based assay using CellTiter-Glo® Luminescent Cell Viability Assay (G7570, Promega, Madison, WI, USA). RAW 264.7 cells and BMMs were seeded in a white opaque-bottom 96-well plate. After treatment with RANKL (30 ng/mL) and E_2_ (10^-8^ M) for 3, 6 and 24 h, 50 μl of culture media was replaced with 50 μl of assay reagent (CellTiter-Glo Buffer + CellTiter-Glo Substrate). Contents were mixed for 2 minutes on an orbital shaker to promote cell lysis, followed by a 10 min incubation at room temperature. The luminescence signal was monitored in a Cytation 3 reader (BioTek Instruments, Winooski, VT, USA).

### Evaluation of mitochondrial membrane potential

To indirectly assess mitochondrial transmembrane potential, RAW 264.7 cells and BMMs were seeded in Corning 96-well black polystyrene microplate and exposed to RANKL (30 ng/mL) and E_2_ (10^-8^ M) for 3 h and 6 h, respectively. After treatment, cells were incubated with assay medium (DMEM D5030 supplemented with 25 mM D-(+)-Glucose, 4 mM L-Glutamine, 1 mM Sodium Pyruvate) containing 100 nM of tetramethylrhodamine (TMRM; T668, Invitrogen, Waltham, MA, USA) for 30 min at 37°C. The fluorescent signal was measured at 548 nm excitation wavelength and 574 nm emission wavelength in a Cytation™ 3 microplate reader (BioTek Instruments Inc., Winooski, VT, USA). As a positive control, 1 μM FCCP was added at the beginning of TMRM incubation. At the end of the assay, the cells were fixed overnight with 10% TCA at 4°C for further SRB normalization (section 2.6).

### Determination of mitochondrial superoxide production

Mitochondrial-specific probe MitoSOX Red Mitochondrial Superoxide Indicator (M36008, Thermo Fisher Scientific, Hampton, NA, USA) was used to evaluate mitochondrial superoxide anion. RAW 264.7 cells and BMMs were seeded in a Corning 96-well black polystyrene microplate and treated with RANKL (30 ng/mL), with or without E_2_ (10^-8^ M) for 3 and 6 h, respectively. After treatment, medium was replaced by the assay medium (described above) containing 5 μM of MitoSOX. Cells were incubated with MitoSOX for 30 min at 37°C. After incubation, the fluorescent signal was measured at a 510 nm excitation wavelength and 580 nm emission wavelength in a Cytation™ 3 microplate reader. A kinetic assay was performed for 120 minutes, with the fluorescence being measured every 2 min. As a positive control, 3 μM antimycin was added at the beginning of MitoSOX incubation. In the end, a linear regression analysis was performed to evaluate the MitoSOX oxidation rate. After fixing with 10% TCA overnight at 4°C, SRB assay (section 2.6) was performed to normalize the data.

### Sulforhodamine B (SRB) assay

The SRB assay was used to evaluate the cellular protein content and it was used in the most assay to normalize data (Vichai & Kirtikara, 2006). After fixation with 10% TCA, cells were incubated with 30 μl of 0.05% (w/w) SRB (S9012, Sigma-Aldrich, St. Louis, MO, USA) in 1% (v/v) acetic acid in methanol for 1 h at room temperature. The unbound SRB was removed by rinsing the cells four times with 1% (v/v) acetic acid. The dye bounded to the cells’ proteins was solubilized with 200 μl of 10 mM Tris-base solution (pH 10.5). SRB absorbance was measured at 510 nm in Cytation 3 reader (BioTek Instruments Inc., USA).

### Complex I activity

RAW 264.7 cells were cultured in 60 mm cell culture dish with RANKL (30 ng/mL) with or without E_2_ (10^-8^ M) for 3 h. Cells were collected, and the pellet was resuspended in 300 μl of phosphate buffer (PB; 50 mM, pH 7.0). The suspension was homogenized by 30 passages through a 27-gauge needle followed by three cycles of freeze/thawing in liquid nitrogen. Soluble protein content was quantified using the Pierce™ BCA Protein Assay Kit (23225, Thermo Fisher Scientific, Hampton, NA, USA), and the samples were diluted to a final concentration of 1 mg/mL. Complex I (nicotinamide adenine dinucleotide (NADH) dehydrogenase) activity was determined according to a previously described protocol (Spinazzi et al., 2012). Briefly, to each well of a Corning 96-well black polystyrene microplate, it was added 130 μl of H_2_O, 21 μl of PB (0.5 M), pH 7.5), 12.6 μl of BSA (50 mg/mL), 6 μl KCN (10 mM), 10 μl NADH (2 mM, N8129, Sigma-Aldrich, St. Louis, MO, USA) and 10 μl of each sample (1 mg/mL). Complex I activity was measured for 5 min, followed by adding 6 μl decylubiquinone (10 mM, D7911, Sigma-Aldrich, St. Louis, MO, USA) to start the reaction. The decrease in absorbance at 340 nm due to oxidation of NADH was followed during 10 min. The maximal activity was determined using the slope of the experimental values’ linear regression and is expressed in enzyme units (U) obtained by the Beer-Lambert law with l = 0.484 cm and L = 6.2 mmol-1.cm-1. NADH dehydrogenase activity of the negative controls was subtracted from their respective samples.

Citrate synthase activity was used to normalize complex I activity. The samples were prepared as described above. Citrate synthase activity was determined according to a previously described protocol (Spinazzi et al., 2012). To each well, 10 μl of each sample (1 mg/mL), 80 μl of 1 mM 5,5’-Dithiobis(2-nitrobenzoic acid) (DTNB), 10 μl of 4 mM acetyl-CoA, 10 μL of 1% Triton X-100 (in 50 mM PB pH 7.0) and 20 μl of H_2_O was added. The baseline activity was measured for 2 min. The reaction was started by adding 20 μl of oxaloacetate (10 mM), and the increase in absorbance due to the consumption of DTNB and formation of 5-Mercapto-2-nitrobenzoic acid (TNB) was followed at 412 nm for 5 min. Each sample was measured in triplicates. Oxaloacetate was replaced by the corresponding volume of 50 mM PB (pH 7.0) for the negative controls. Citrate synthase activity was determined using the slope experimental values’ linear regression and was expressed in enzyme units (U) obtained by the Beer-Lambert law with l = 0.346 cm and L = 13.6 mmol1.cm-1. Citrate synthase activity of the negative controls was subtracted from their respective samples

### Mitochondrial DNA (mtDNA) copy number measurements

mtDNA copy number was determined using quantitative polymerase chain reaction (qPCR). RAW 264.7 cells were seeded in 6-well plate and treated with RANKL (30 ng/mL) and E_2_ (10^-8^ M) for 3 h. After treatment, cells were collected by scraping and washed with ice-cold 1x PBS. DNA was isolated using the Quiagen DNeasy kit (69104, Qiagen, Hilden, Germany) according to the manufacturer’s instructions. Total DNA was quantified at 260 nm absorbance using NanoDrop 2000 (ThermoScientific, Waltham, MA, USA). qPCR was performed using SsoFast™ EvaGreen Supermix (1725204, Bio-Rad, Hercules, CA, USA) in a CFX96 real time-PCR system (Bio-Rad, Hercules, CA, USA) and the DNA copy number was calculated based on the amplification of *mito1* (encoded on the mitochondrial genome, presents a variable quantity in each cell) and beta-2-microglobulin (*b2m*; encoded on the nuclear genome, presents a fixed quantity in each cell) as s detailed in Malik et al. (Malik et al., 2016). Mouse primers used for mito1 were forward 5’-CTAGAAACCCCGAAACCAAA-3’ and reverse 5’-CCAGCTATCACCAAGCTCGT-3’; for b2m were forward 5’-ATGGGAAGCCGAACATACTG-3’ and reverse 5’-CAGTCTCAGTGGGGGTGAAT -3’. Total DNA (25 ng) was loaded in Hard-Shell 96-well PCR plate, and the amplification started with an initial cycle of 2 min at 98°C for initial denaturation, followed by 40 cycles of 5 sec at 98°C plus 5 sec at 60°C and 5 sec at 65°C. EvaGreen fluorescence was recorded at the end of each cycle to determine Cq. Several dilutions of the control samples and DNA-free water for each primer were used to assess amplification efficiency and exclude contamination. Each reaction was performed in triplicate and presented an efficiency between 95 and 105%. Mitochondrial copy number was estimated by the ratio between the number of copies of *mito1* template divided by the number of copies of *b2m* template. The normalized expression was calculated by the comparative quantification algorithm ΔΔCt, using CFX96 Manager software (v. 3.0; Bio-Rad, Hercules, CA, USA) (RRID:SCR_017251).

### Quantitative real-time polymerase chain reaction: qRT-PCR

RAW 264.7 cells were seeded in 6-well tissue culture plates and treated with RANKL (30 ng/mL) and E_2_ (10^-8^ M) during 3 h. TripleXtractor (GB23, GRiSP, Porto, Portugal) was used to isolate RNA fractions from whole samples. Briefly, after treatments, the media was removed and 1 mL of TripleXtractor was added to each well. The reagent was mixed with the cellular extracts and incubated at room temperature for 5 min to disrupt nucleoprotein complexes. Then, 200 μl of chloroform (650498, Sigma-Aldrich, St. Louis, MO, USA) was added and mixed vigorously for 15 sec. The samples were incubated for 5 min at room temperature and centrifuged at 12000 x g for 15 min at 4°C. This process resulted in a separation into 3 phases: aqueous phase (RNA), interphase (DNA) and organic phase (protein). The aqueous phase was transferred to a new microtube, and total RNA was extracted with RNeasy mini kit (74104, Qiagen) following the manufacture’s protocol. Total RNA was quantified at 260 nm absorbance using NanoDrop 2000 (Thermo Fisher Scientific, Hampton, New Hampshire, USA). Then, 2 μg of RNA template was converted to cDNA using iScript™ cDNA Synthesis Kit (1708891, Bio-Rad, Hercules, CA, USA), according to the manufactures’ instructions. RT-PCR in a CFX96 real time-PCR system (Bio-Rad, Hercules, CA, USA), using SsoFast™ EvaGreen Supermix and the primers defined in table 1.

**Table 1.**
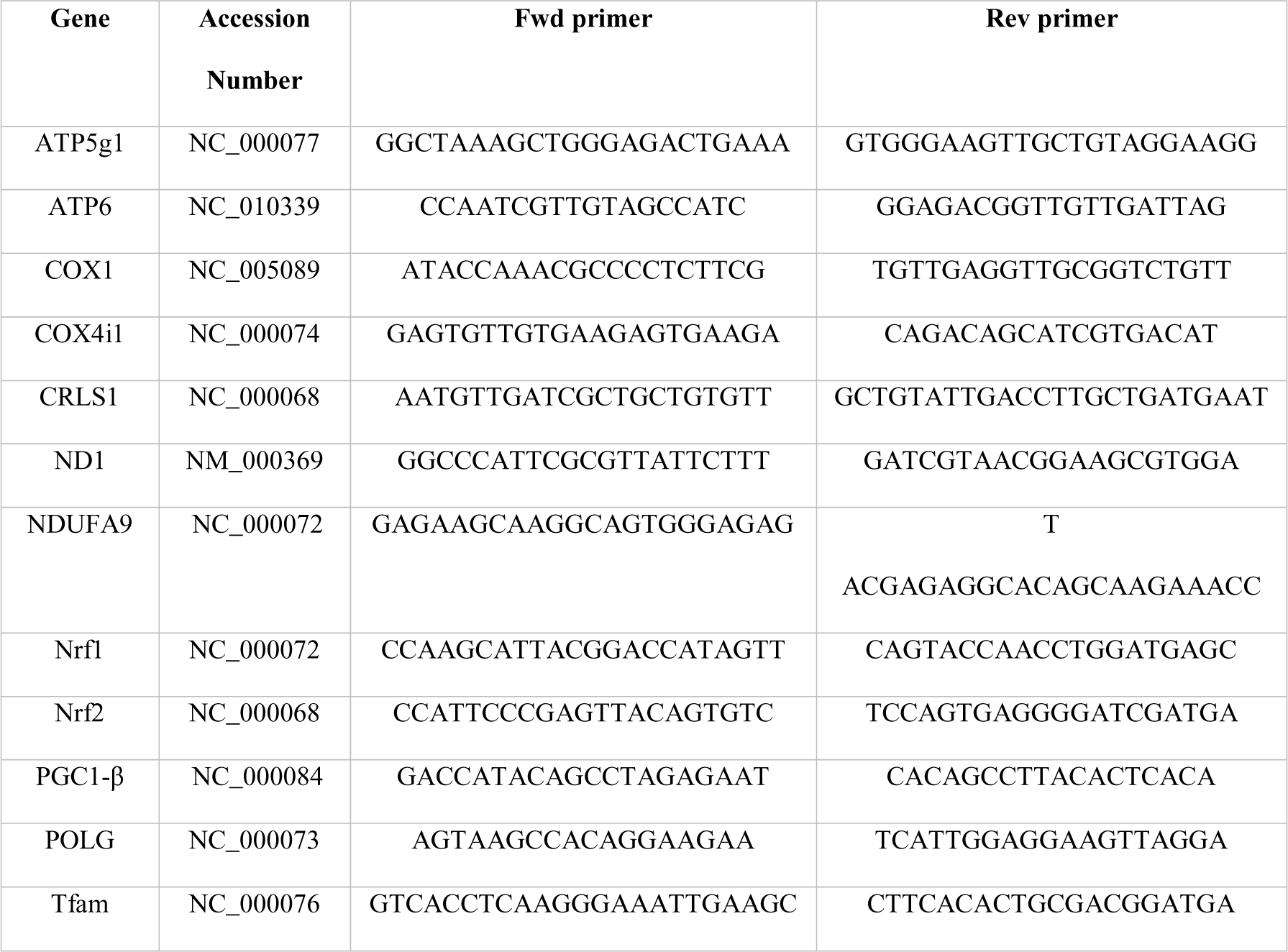
List of forward (Fwd) and reverse (Rev) primers used for qRT-PCR, respective gene and accession number

Amplification of 25 ng of cDNA was performed with an initial cycle of 30 sec at 95°C for initial denaturation, followed by 40 cycles of 5 sec at 95°C plus 5 sec at 60°C. EvaGreen fluorescence was recorded at the end of each cycle to determine Cq. Several dilutions of the control samples and DNA-free water for each primer were used to assess amplification efficiency and exclude contamination. Each reaction was performed in triplicate and presented an efficiency between 95 and 105%. Relative normalized expression was determined by the CFX96 Manager software (v. 3.0; Bio-Rad, Hercules, CA, USA) (RRID:SCR_017251), using b2m and β-actin as reference genes.

### Cardiolipin content evaluation

To measure cardiolipin content in cell lysates, the Cardiolipin Assay kit (MAK362, Sigma-Aldrich, St. Louis, MO, USA) was used according to the manufacturer’s instructions. Briefly, RAW 264.7 cells were cultured in 60 mm cell culture dish with RANKL (30 ng/mL) with or without E_2_ (10^-8^ M) for 6 h. Cells were collected by scraping and centrifuged at 300 x g for 5 min. The pellet was resuspended in 75 μl of CL Assay buffer and submitted to three cycles of freeze/thawing. The lysate was centrifuged at 10000 x g for 4 min at 4°C, and protein was quantified using the Pierce™ BCA Protein Assay Kit, using BSA as a standard. To measure cardiolipin content, 10 μg of protein and 50 μl of reaction mix was added to a 96-wells white opaque-bottom plate. After 10 min incubation at room temperature, the fluorescence was measured at 340 nm excitation wavelength and 480 nm emission wavelength in a Cytation™ 3 microplate reader (BioTek Instruments Inc., Winooski, VT, USA).

### Mitochondrial metabolic phenotyping

MitoPlate S-1 plates from Biolog (Biolog, Hayward, CA, USA) (Bochner et al., 2011) were used according to manufacturer’s instructions. Each substrate was present in one well, and the mitochondrial substrate oxidizing capacity was measured by evaluating the rates of electron flow from the substrates to the electron transport chain, where a tetrazolium redox dye acts as an electron acceptor and turns purple after reduction. Briefly, RAW 264.7 cells and BMMs were seeded in 60 mm cell culture dishes and treated with RANKL (30 ng/mL) and E_2_ (10^-8^) for 24 h. The cells were harvested, washed twice with 1x PBS, and suspended in 1 mL of 1x Biolog Mitochondrial Assay Solution (BMAS). The suspension was filtered through a 70 μm nylon strainer. Per each well of the MitoPlate, 30 μl of cell suspension containing 7x10^4^ cells for RAW 264.7 or 15x10^4^ cells for BMMs, 30 μl of reaction mix (15 μl BMAS, 10 μl Redox Dye MC, 2.5 μl saponin (75 μg/mL; SAE0073, Sigma-Aldrich, St. Louis, MO, USA) and 2.5 μl sterile water was added and incubated for 1h at 37°C. The MitoPlate was inserted in the Omnilog (Biolog, Hayward, CA, USA) set at 37°C and the reduction of the tetrazolium redox dye MC was kinetically measured for 24 h, with 5 min intervals. The initial rate was calculated using the Biolog software data analysis (v. 1.7).

### Hoechst 33342 staining assay

RAW 264.7 were seeded in Corning 96-well black polystyrene microplate and treated with RANKL (30 ng/mL) and E_2_ (10^-8^ M) for 3, 6, 12 and 24 h. After treatment, cells were rinsed fixed with 4% paraformaldehyde (PFA) for 15 min at room temperature. The nuclei were stained with 1 μg/mL Hoechst 33342 solution in 1x PBS for 15 min at room temperature. Nuclei counting was performed using an INCell Analyzer 2200 (GE Healthcare, Chicago, IL, USA) cell imaging system. Images were acquired using a 10x objective (Nikon 10x/0.45, Plan Apo, CFI/60) using the DAPI channel (400-418/478-498 nm) and analyzed using the INCell Analyzer 1000 analysis software - Developer Toolbox (RRID:SCR_015790). The representative images were processed using TotalLab TL120 Software (version 2009).

### Measurement of caspase 3 and 9-like activities

RAW 264.7 cells were seeded in 60 mm cell culture dish and treated with RANKL (30 ng/mL) and E_2_ (10^-8^) for 6, 12 and 24 h. Total cellular extracts were obtained after two centrifugation steps at 1000 x g for 5 min at 4°C. Cellular pellets were resuspended in 100 μL of cell lysis buffer containing 50 mM HEPES (pH 7.4, 100 mM NaCl, 0.1% 3[(3-cholamidopropyl)dimethy-lammonio]-propanesulfonic acid (CHAPS), 0.1 mM EDTA and supplemented with 100 μM of phenylmethylsulphonyl fluoride (PMSF) and 2 mM of dithiothreitol (DTT). The suspension was homogenized by 30 passages through a 27-gauge needle, followed by three cycles of freeze/thawing in liquid nitrogen. Protein content was determined by the Bradford method (Bradford, 1976), using bovine serum albumin (BSA) as standard. To measure caspase 3 and caspase 9-like activity, 25 μg or 50 μg of total protein were used, respectively, and incubated in a reaction buffer containing 10 mM DTT, 25 mM HEPES (pH 7.4), 10% sucrose, 0.1% CHAPS and 100 μM caspase substrate (Ac-DEVD-p-nitroanilide (pNA) (ALX-260-03, Enzo Life Science, Farmingdale, NY, USA) for caspase 3 or Ac-LEHD-pNA (ALX-260-081, Enzo Life Science) for caspase 9, for 2 h at 37 °C. Caspase-like activities were determined by following the cleavage of the chromophore pNA from the substrate. pNA absorbance was measured at 405 nm in Cytation 3 reader (BioTek Instruments Inc., Winooski, VT, USA). A pNA (N2128, Sigma-Aldrich, St. Louis, MO, USA) standard curve was used to calibrate the method.

### Nutlin-3a treatment

RAW 264.7 cells and BMMs were seeded in 60 mm cell culture dishes and treated with E_2_ (10^-8^ M), RANKL (30 ng/mL) plus E_2_, or Nutlin-3a (SML0580, Sigma-Aldrich, St. Louis, MO, USA). for 12 or 24 h, respectively. P53 protein total levels and pSer392-p53 were determined by western blotting as described below (2.17). Caspase 3 activity was determined as described above (2.14).

### Subcellular fractionation

RAW 264.7 cells were cultured in 60 mm cell culture dish with RANKL (30 ng/mL) with or without E_2_ (10^-8^ M) during 3 or 6 h. Cells were washed with ice-cold PBS and the pellet was lysed with extraction buffer (50 mM Tris-HCl (pH 8.0), 2 mM EDTA, 0.5% Nonidet P-40, 20% glycerol, 25 mM β-glycerophosphate and 0.5 mM PMSF for 5 min on ice, and then centrifuged for 5 min at 720 x g and 4°C. Supernatants were used as cytosolic extracts. For mitochondrial purification, the pellet was resuspended in 100 μl of NKM buffer (1 mM Tris-HCl (pH 7.4), 0.13 M NaCl, 5 mM KCl and 7.5 mM MgCl_2_). The suspension was centrifuged at 370 x g for 10 min and resuspended in NKM buffer twice. After these washing steps, the cells were resuspended in 60 μl of homogenization buffer (10 mM Tris-HCl (pH 6.7), 10 mM KCl, 0.15 mM MgCl_2_, 1 mM PMSF and 1 mM DTT) and incubated on ice for 10 min. The suspension was homogenized by 30 passages through a 27-gauge needle, and the homogenate was transferred to a new centrifuge tube containing 10 μl of 2 M sucrose and mixed gently. Intact cells, nuclei, and other debris were pelleted by centrifuging the suspension for 5 min at 1200 x g. The supernatant was transferred to a new microtube, and this process was repeated twice. The suspension was centrifuged at 7000 x g for 10 min. The pellet was used as the mitochondria-enriched fraction and was resuspended in 30 μl of mitochondrial suspension buffer (10 mM Tris-HCl (pH 6.7), 0.15 mM MgCl_2_, 0.25 mM sucrose, 1 mM PMSF and 1 mM DTT). Both cytosolic and mitochondrial extracts were quantified using the Pierce™ BCA Protein Assay Kit, using BSA as a standard, and were used for western blotting analysis.

### Western blotting analysis

RAW 264.7 cells were seeded in 6-well cell culture plates and treated with RANKL (30 ng/mL) and E_2_ (10^-8^ M) for 3 h. After treatment, cells washed with PBS and pellet was resuspended in 100 μl of RIPA buffer (50 mM Tris pH 8, 150 mM NaCl, 5 mM EDTA, 15 mM MgCl_2_ and 1% TritonX-100 supplemented with protease inhibitors (protease inhibitor cocktail (PIC), 0.5 mM PMSF, 20 mM NaF, 400 mM NaM, 500 mM sodium butyrate and 0.5% DOC. The suspension was maintained on ice for 1 h and, vortexed every 10 min. Soluble protein content was quantified using the Pierce™ BCA Protein Assay Kit, using BSA as a standard. The samples were diluted to a final concentration of 2 mg/mL with Laemelli sample buffer (188 mM Tris-HCl, 218 mM SDS, 8% glycerol, and 500 μM Bromophenol Blue (B5525, Sigma-Aldrich, St. Louis, MO, USA). An equal amount of protein, 30 μg, was separated by electrophoresis on a 10-14% acrylamide/bis-acrylamide gel and transferred to a PVDF membrane. Membranes were stained with Ponceau S reagent to confirm equal protein loading and blocked with 5% BSA for 2 h at room temperature with constant shaking. After blocking, membranes were incubated overnight at 4°C with antibodies directed against Membrane Integrity cocktail (1:1000, ab110414, Abcam, Cambridge, UK), p-p53 (1:500, sc-51690, Santa Cruz Biotechnology, Dallas, TX, USA), Bak (1:1000, sc-518146, Santa Cruz Biotechnology, Dallas, TX, USA), Bax (1:1000, 2772, Cell Signaling, Danvers, MA, USA), denaturated form of OXPHOS complexes cocktail (1:1000, MS604, MitoSciences, Eugene, OR, USA), Estrogen receptor alpha (ERα; 1:500, ab16460, Abcam, Cambridge, UK), GAPDH (1:1000, sc-59540, Santa Cruz Biotechnology, Dallas, TX, USA), TOM 20 (1:1000, sc-11415, Santa Cruz Biotechnology, Dallas, TX, USA) and β-actin (1:5000, MAB1501, Chemicon International, Thermo Fisher Scientific, Hampton, New Hampshire, USA). Membranes were then incubated with horseradish peroxidase (HRP)-conjugated goat anti-mouse IgG (1:5000, CS7076, Cell Signaling, Danvers, MA, USA) and goat anti-rabbit IgG (1:5000, CS7074, Cell Signaling, Danvers, MA, USA) secondary antibodies for 1 h at room temperature. For the chemiluminescence detection, membranes were incubated with the Clarity Western ECL Substrate (1705061, Bio-Rad, Hercules, CA, USA) and incubated for 5 min at low light conditions. The chemiluminescence was detected by Imager Imaging System (VWR, PA, USA). Band density was measured by using TotalLab TL120 Software (version 2009).

### Computational data analysis

Data analysis comprised the training of supervised decision trees to identify the inhibitory effect of E_2_ on the RANKL-induced mitochondrial activity. The RAW 264.7 dataset contains 12 RANKL samples and 12 RANKL plus E_2_ samples, whereas the BMMs dataset contains 18 RANKL samples and 18 RANKL plus E_2_ samples Six cellular oxygen consumption measurements were considered in the study: {ATP-linked OCR, Basal Respiration, Maximal Respiration, Non-mitochondrial oxygen consumption, Proton Leak and Spare Respiratory Capacity}. The computational analysis of the data was performed using Python 3, version 3.7.4. We relied on the Pandas package (McKinney, 2010) to load, store and process the data. The scikit-learn library (Pedregosa et al., 2011) was used to train the decision trees. Given the moderate sample sizes, a stratified cross-validation strategy, with 4 folds, was adopted to fit the trees. Hyper-parameters were tuned using the RandomizedSearchCV method from scikit-learn, an efficient grid search variant. The selected settings are the following: {minimum number of samples to split a node: 10, 5, respectively for the information from the cellular line and the primary bone marrow cells; number of features to consider when splitting a node: square root of the total number of features; maximum tree depth: 4; split criterion: gini/entropy, respectively for the information from the cellular line and the primary bone marrow cells}.

Radar charts were used to display multivariate data in the form of a two-dimensional chart of quantitative variables. The average value of the samples was considered for each pair condition/substrate. Radar charts were created with the Plotly graphing library (Plotly Technologies Inc., 2015), whereas Matplotlib (Hunter, 2007) was used to create the remaining figures.

### Statistical Analysis

Statistical analysis and graphical design were performed in GraphPad Prism 8 software (Graphpad Software, CA, USA). Data are represented by mean values with standard error of the mean. Statistical significance between the three conditions in the different timepoints, was analyzed by two-way ANOVA following

Tukey’s multiple comparisons test. Additionally, when comparing the different conditions in a single timepoint, it was performed one-way ANOVA analysis with Tukey’s multiple comparisons test.

## Author contribution

A.M.C generated and analyzed the data. B.S. and F.B.P performed the computational analysis of the data. A.M.C wrote the manuscript, KAN, M.A., P.J.O. and V.A.S supervised the research. All authors reviewed the manuscript.

## Acknowledgements

This work was funded by the European Regional Development Fund (ERDF), through Operational Programme for Competitiveness and Internationalization (COMPETE) and Portuguese funds national funds via *Fundação para a Ciência e Tecnologia* (FCT), under projects: IF/01182/2015 and POCI-01-0145-FEDER-029297, AMC (SFRH/BD/140817/2018) was supported by FCT PhD fellowship; VAS also was supported by Research Executive Agency (REA) (’the Agency’) by the European Commission, under the project MIA-Portugal, Grant Agreement number: 85752

---

**Supplementary Figure S1.**
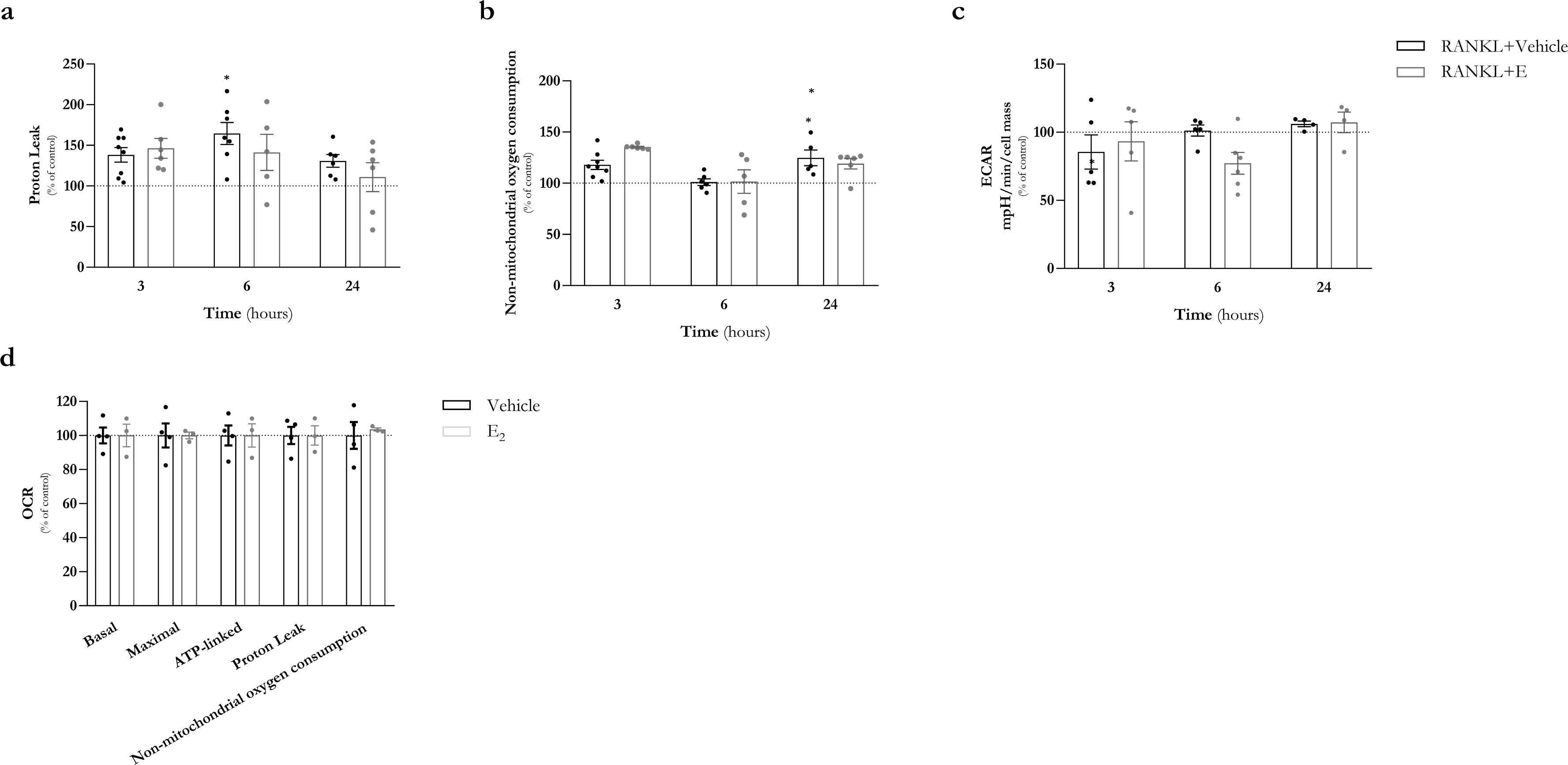
Time-dependent effects of RANKL and E_2_ in oxygen consumption, proton leak, non-mitochondrial oxygen consumption and extracelular acidification in RAW 264.7 cells. RAW 264.7 cells were treated RANKL (30 ng/mL) with or without E_2_ (10^-8^ M) for 3, 6 and 24 h. (A) Proton leak, (B) Non-mitochondrial oxygen consumption and (C) Extracellular acidification rate (ECAR) parameters were evaluated using the Mitostress Seahorse assay. (D) Comparison of oxygen consumption rate (OCR) parameters between control and E_2_ treatment, without RANKL. The results were normalized for the control condition (cells without treatment, 100% marked by a doted line). Control vs RANKL (black) and RANKL vs RANKL+E_2_ (grey) significance was accepted with p<0.05, using two-way ANOVA analysis with Tukey’s multiple comparisons test.

**Supplementary Figure S2.**
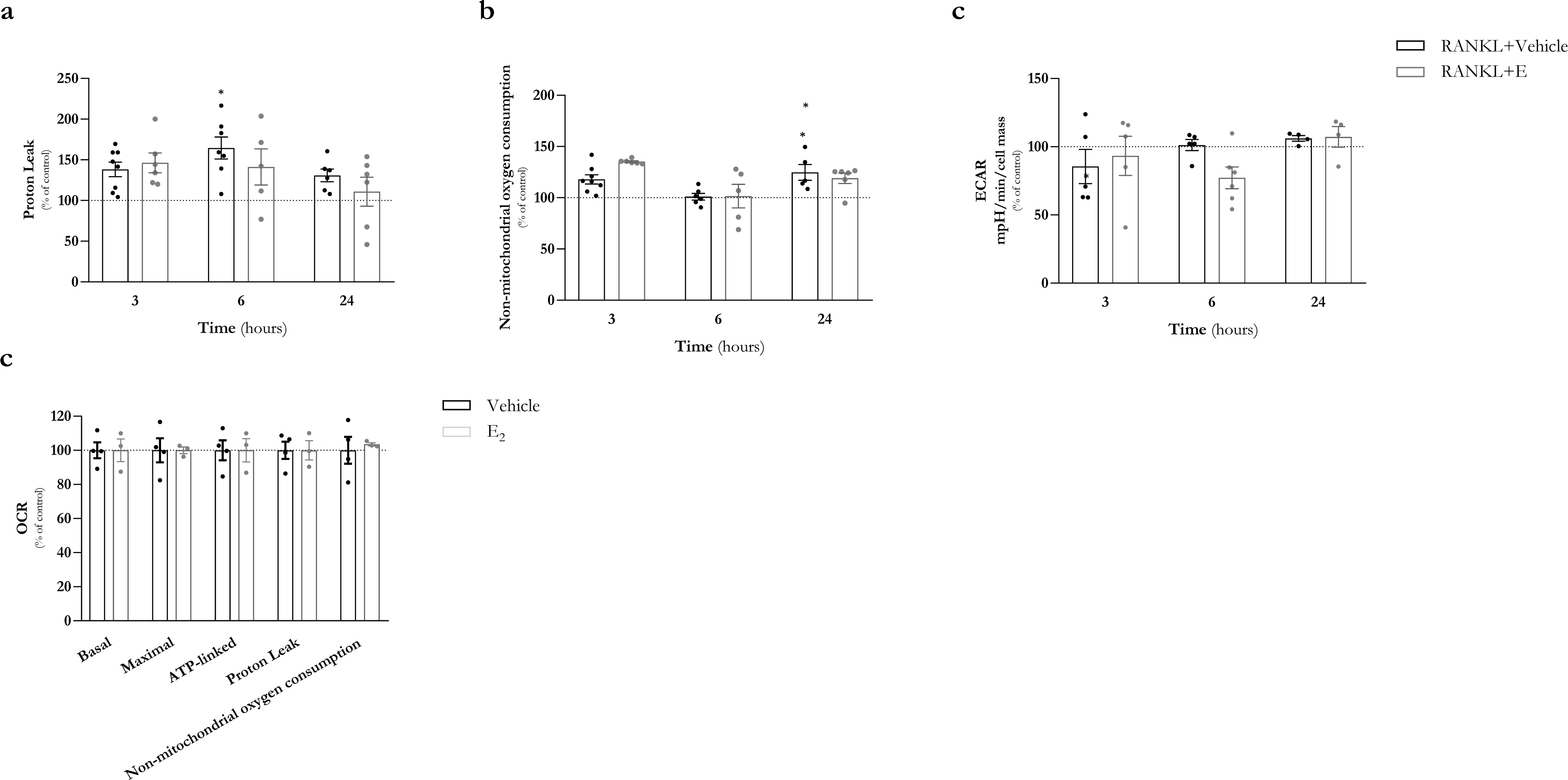
Time-dependent effects of RANKL and E_2_ in oxygen consumption, proton leak, non-mitochondrial oxygen consumption and extracelular acidification in BMMs. BMMs were suplemented with M-CSF (30 ng/mL) and treated RANKL (30 ng/mL) with or without E_2_ (10^-8^ M) for 6 and 24 h. **(A)** Proton leak, **(B)** Non-mitochondrial oxygen consumption and **(C)** Extracellular acidification rate (ECAR) parameters were evaluated using the Mitostress Seahorse assay. **(D)** Comparison of oxygen consumption rate (OCR) parameters between control and E_2_ treatment, without RANKL. The results were normalized for the control condition (cells with M-CSF, 100% marked by a doted line). M-CSF vs RANKL (black) and RANKL vs RANKL+E_2_ (grey) significance was accepted with p<0.05, using two-way ANOVA analysis with Tukey’s multiple comparisons test.

**Supplementary Figure S3.**
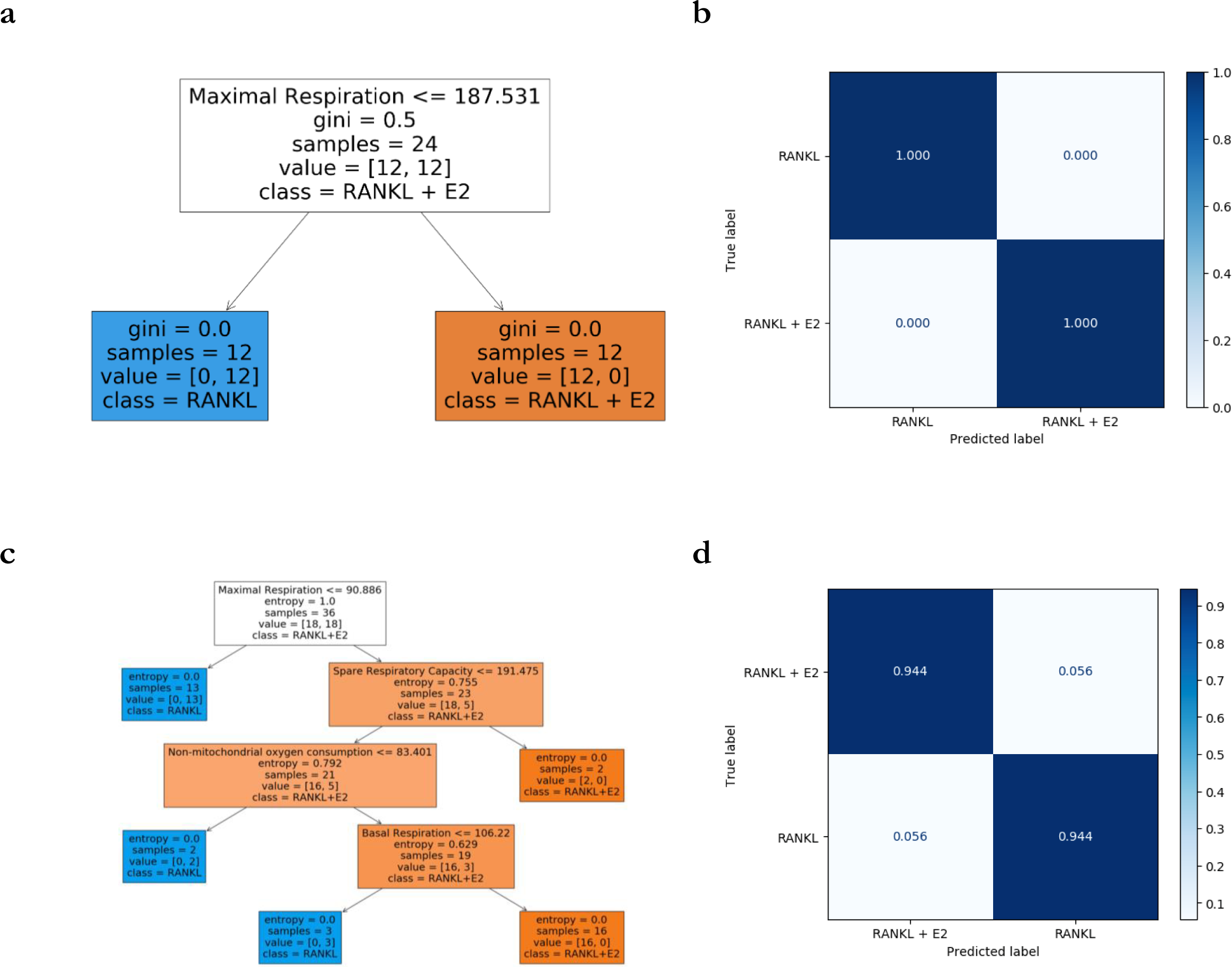
A computational supervised classifier using OCR measurements confirms a clear effect of E_2_ on the RANKL treatment in RAW 264.7 cells and BMMs. Decision trees were trained with cellular oxygen consumption measurements for both RAW 264.7 cells and BMMs. The RAW 264.7 dataset contains 15 RANKL samples and 16 RANKL plus E_2_ samples. The corresponding tree model (A) perfectly separates the classes, as confirmed by the confusion matrix obtained with test data (B). The BMMs dataset contains 18 RANKL samples and 18 RANKL plus E_2_ samples. The model trained with primary cells (C) achieves 95% test accuracy (D).

## Bibliography

Almeida, M., Laurent, M. R., Dubois, V., Claessens, F., O’Brien, C. A., Bouillon, R., Vanderschueren, D., & Manolagas, S. C. (2017). Estrogens and androgens in skeletal physiology and pathophysiology. Physiological Reviews, 97(1), 135–187. https://doi.org/10.1152/physrev.00033.2015

An, E., Narayanan, M., Manes, N. P., & Nita-Lazar, A. (2014). Characterization of functional reprogramming during osteoclast development using quantitative proteomics and mRNA profiling. Molecular and Cellular Proteomics, 13(10), 2687–2704. https://doi.org/10.1074/mcp.M113.034371

Arnett, T. R., & Orriss, I. R. (2018). Metabolic properties of the osteoclast. In Bone (Vol. 115, pp. 25–30). Elsevier Inc. https://doi.org/10.1016/j.bone.2017.12.021

Bauer, T. M., & Murphy, E. (2020). Role of mitochondrial calcium and the permeability transition pore in regulating cell death. In Circulation Research (pp. 280–293). Lippincott Williams and Wilkins. https://doi.org/10.1161/CIRCRESAHA.119.316306

Berger, C. E., Qian, Y., Liu, G., Chen, H., & Chen, X. (2012). p53, a target of estrogen receptor (ER) α, modulates DNA damage-induced growth suppression in ER-positive breast cancer cells. Journal of Biological Chemistry, 287(36), 30117–30127. https://doi.org/10.1074/jbc.M112.367326

Bochner, B. R., Siri, M., Huang, R. H., Noble, S., Lei, X. H., Clemons, P. A., & Wagner, B. K. (2011). Assay of the multiple energy-producing pathways of mammalian cells. PLoS ONE, 6(3). https://doi.org/10.1371/journal.pone.0018147

Castrogiovanni, C., Waterschoot, B., de Backer, O., & Dumont, P. (2018). Serine 392 phosphorylation modulates p53 mitochondrial translocation and transcriptionindependent apoptosis. Cell Death and Differentiation, 25(1), 190–203. https://doi.org/10.1038/cdd.2017.143

Da, W., Tao, L., & Zhu, Y. (2021). The Role of Osteoclast Energy Metabolism in the Occurrence and Development of Osteoporosis. In Frontiers in Endocrinology (Vol. 12). Frontiers Media S.A. https://doi.org/10.3389/fendo.2021.675385

Dewson, G., Ma, S., Frederick, P., Hockings, C., Tan, I., Kratina, T., & Kluck, R. M. (2012). Bax dimerizes via a symmetric BH3:groove interface during apoptosis. Cell Death and Differentiation, 19(4), 661–670. https://doi.org/10.1038/cdd.2011.138

Dougall, W. C., Glaccum, M., Charrier, K., Rohrbach, K., Brasel, K., de Smedt, T., Daro, E., Smith, J., Tometsko, M. E., Maliszewski, C. R., Armstrong, A., Shen, V., Bain, S., Cosman, D., Anderson, D., Morrissey, P. J., Peschon, J. J., & Schuh, J. A. (1999). RANK is essential for osteoclast and lymph node development. Genes and Development, 13(18), 2412–2424. https://doi.org/10.1101/gad.13.18.2412

Dudek, J. (2017). Role of cardiolipin in mitochondrial signaling pathways. In Frontiers in Cell and Developmental Biology (Vol. 5, Issue SEP). Frontiers Media S.A. https://doi.org/10.3389/fcell.2017.00090

Green, D. R., & Kroemer, G. (2009). Cytoplasmic functions of the tumour suppressor p53. Nature, 458(7242), 1127–1130. https://doi.org/10.1038/nature07986

Gureev, A. P., Shaforostova, E. A., & Popov, V. N. (2019). Regulation of mitochondrial biogenesis as a way for active longevity: Interaction between the Nrf2 and PGC-1α signaling pathways. In Frontiers in Genetics (Vol. 10, Issue MAY). Frontiers Media S.A. https://doi.org/10.3389/fgene.2019.00435

Hsu, Y.-T., Wolter, K. G., & Youle, R. J. (1997). Cytosol-to-membrane redistribution of Bax and Bcl-X L during apoptosis (programmed cell deathBcl-2Baxdiphtheria toxin) (Vol. 94). www.pnas.org.

Hughes, D. E., Dai, A., Tiffee, J. C., Li, H. H., Munoy, G. R., & Boyce, B. F. (1996). Estrogen promotes apoptosis of murine osteoclasts mediated by TGF-. Nature Medicine, 2(10), 1132–1135. https://doi.org/10.1038/nm1096-1132

Indo, Y., Takeshita, S., Ishii, K. A., Hoshii, T., Aburatani, H., Hirao, A., & Ikeda, K. (2013). Metabolic regulation of osteoclast differentiation and function. Journal of Bone and Mineral Research, 28(11), 2392–2399. https://doi.org/10.1002/jbmr.1976

Ishii, K. A., Fumoto, T., Iwai, K., Takeshita, S., Ito, M., Shimohata, N., Aburatani, H., Taketani, S., Lelliott, C. J., Vidal-Puig, A., & Ikeda, K. (2009). Coordination of PGC-1β and iron uptake in mitochondrial biogenesis and osteoclast activation. Nature Medicine, 15(3), 259–266. https://doi.org/10.1038/nm.1910

Kim, H. N., Ponte, F., Nookaew, I., Ucer Ozgurel, S., Marques-Carvalho, A., Iyer, S., Warren, A., Aykin-Burns, N., Krager, K., Sardao, V. A., Han, L., de Cabo, R., Zhao, H., Jilka, R. L., Manolagas, S. C., & Almeida, M. (2020). Estrogens decrease osteoclast number by attenuating mitochondria oxidative phosphorylation and ATP production in early osteoclast precursors. Scientific Reports, 10(1), 1–17. https://doi.org/10.1038/s41598-020-68890-7

Kitagawa, M., Aonuma, M., Lee, S. H., Fukutake, S., & McCormick, F. (2008). E2F-1 transcriptional activity is a critical determinant of Mdm2 antagonist-induced apoptosis in human tumor cell lines. Oncogene, 27(40), 5303–5314. https://doi.org/10.1038/onc.2008.164

Kracikova, M., Akiri, G., George, A., Sachidanandam, R., & Aaronson, S. A. (2013). A threshold mechanism mediates p53 cell fate decision between growth arrest and apoptosis. Cell Death and Differentiation, 20(4), 576–588. https://doi.org/10.1038/cdd.2012.155

Kruiswijk, F., Labuschagne, C. F., & Vousden, K. H. (2015). P53 in survival, death and metabolic health: A lifeguard with a licence to kill. Nature Reviews Molecular Cell Biology, 16(7), 393–405. https://doi.org/10.1038/nrm4007

Kuznetsov, A. v., Margreiter, R., Amberger, A., Saks, V., & Grimm, M. (2011). Changes in mitochondrial redox state, membrane potential and calcium precede mitochondrial dysfunction in doxorubicin-induced cell death. Biochimica et Biophysica Acta - Molecular Cell Research, 1813(6), 1144–1152. https://doi.org/10.1016/j.bbamcr.2011.03.002

Lemma, S., Sboarina, M., Porporato, P. E., Zini, N., Sonveaux, P., di Pompo, G., Baldini, N., & Avnet, S. (2016). Energy metabolism in osteoclast formation and activity. International Journal of Biochemistry and Cell Biology, 79, 168–180. https://doi.org/10.1016/j.biocel.2016.08.034

Madan, E., Gogna, R., Keppler, B., & Pati, U. (2013). p53 Increases Intra-Cellular Calcium Release by Transcriptional Regulation of Calcium Channel TRPC6 in GaQ3-Treated Cancer Cells. PLoS ONE, 8(8). https://doi.org/10.1371/journal.pone.0071016

Malik, A. N., Czajka, A., & Cunningham, P. (2016). Accurate quantification of mouse mitochondrial DNA without co-amplification of nuclear mitochondrial insertion sequences. Mitochondrion, 29, 59–64. https://doi.org/10.1016/j.mito.2016.05.003

Park-Min, K. H. (2019). Metabolic reprogramming in osteoclasts. Seminars in Immunopathology, 41(5), 565–572. https://doi.org/10.1007/s00281-019-00757-0

Patsch, J. M., Burghardt, A. J., Yap, S. P., Baum, T., Schwartz, A. v., Joseph, G. B., & Link, T. M. (2013). Increased cortical porosity in type 2 diabetic postmenopausal women with fragility fractures. Journal of Bone and Mineral Research, 28(2), 313–324. https://doi.org/10.1002/jbmr.1763

Richards, J. B., Rivadeneira, F., Inouye, M., Pastinen, T. M., Soranzo, N., Wilson, S. G., Andrew, T., Falchi, M., Gwilliam, R., Ahmadi, K. R., Valdes, A. M., Arp, P., Whittaker, P., Verlaan, D. J., Jhamai, M., Kumanduri, V., Moorhouse, M., van Meurs, J. B., Hofman, A.,… Spector, T. D. (2008). Bone mineral density, osteoporosis, and osteoporotic fractures: a genome-wide association study. Www.Thelancet.Com, 371. https://doi.org/10.1016/S0140

Riley, T., Sontag, E., Chen, P., & Levine, A. (2008). Transcriptional control of human p53-regulated genes. Nature Reviews Molecular Cell Biology, 9(5), 402–412.

Rizzotto, D., Zaccara, S., Rossi, A., Galbraith, M. D., Andrysik, Z., Pandey, A., Sullivan, K. D., Quattrone, A., Espinosa, J. M., Dassi, E., & Inga, A. (2020). Nutlin-Induced Apoptosis Is Specified by a Translation Program Regulated by PCBP2 and DHX30. Cell Reports, 30(13), 4355–4369.e6.

Sardão, V. A., Oliveira, P. J., Holy, J., Oliveira, C. R., & Wallace, K. B. (2009). Doxorubicin-induced mitochondrial dysfunction is secondary to nuclear p53 activation in H9c2 cardiomyoblasts. Cancer Chemotherapy and Pharmacology, 64(4), 811–827.

Shao, D., Liu, Y., Liu, X., Zhu, L., Cui, Y., Cui, A., Qiao, A., Kong, X., Liu, Y., Chen, Q., Gupta, N., Fang, F., & Chang, Y. (2010). PGC-1β-Regulated mitochondrial biogenesis and function in myotubes is mediated by NRF-1 and ERRα. Mitochondrion, 10(5), 516–527. https://doi.org/10.1016/j.mito.2010.05.012, M., Casarin, A., Pertegato, V., Salviati, L., & Angelini, C. (2012). Assessment of mitochondrial respiratory chain enzymatic activities on tissues and cultured cells. Nature Protocols, 7(6), 1235–1246.

Stavros C. Manolagas. (2000). Birth and death of bone cells: basic regulatory mechanisms and implications for the pathogenesis and treatment of osteoporosis. Endocrine Reviews, 21(2), 115–137.

Taciak, B., Białasek, M., Braniewska, A., Sas, Z., Sawicka, P., Kiraga, Ł., Rygiel, T., & Król, M. (2018). Evaluation of phenotypic and functional stability of RAW 264.7 cell line through serial passages. PLoS ONE, 13(6), 1–13. https://doi.org/10.1371/journal.pone.0198943

Takamatsu, C., Umeda, S., Ohsato, T., Ohno, T., Abe, Y., Fukuoh, A., Shinagawa, H., Hamasaki, N., & Kang, D. (2002). Regulation of mitochondrial D-loops by transcription factor A and single-stranded DNA-binding protein. EMBO Reports.

Taubmann, J., Krishnacoumar, B., Böhm, C., Faas, M., Müller, D. I. H., Adam, S., Stoll, C., Böttcher, M., Mougiakakos, D., Sonnewald, U., Hofmann, J., Schett, G., Krönke, G., & Scholtysek, C. (2020). Metabolic reprogramming of osteoclasts represents a therapeutic target during the treatment of osteoporosis. Scientific Reports, 10(1).

Teitelbaum, S. L., & Ross, F. P. (2003). Genetic regulation of osteoclast development and function. Nature Reviews Genetics, 4(8), 638–649.

Tovar, C., Rosinski, J., Filipovic, Z., Higgins, B., Kolinsky, K., Hilton, H., Zhao, X., Vu, B. T., Qing, W., Packman, K., Myklebost, O., Heimbrook, D. C., & Vassilev, L. T. (2006). Small-molecule MDM2 antagonists reveal aberrant p53 signaling in cancer: Implications for therapy.

Tsukasaki, M., Huynh, N. C. N., Okamoto, K., Muro, R., Terashima, A., Kurikawa, Y., Komatsu, N., Pluemsakunthai, W., Nitta, T., Abe, T., Kiyonari, H., Okamura, T., Sakai, M., Matsukawa, T., Matsumoto, M., Kobayashi, Y., Penninger, J. M., & Takayanagi, H. (2020). Stepwise cell fate decision pathways during osteoclastogenesis at single-cell resolution. Nature Metabolism, 2(12), 1382–1390.

Vichai, V., & Kirtikara, K. (2006). Sulforhodamine B colorimetric assay for cytotoxicity screening. Nature Protocols, 1(3), 1112–1116.

Zeng, R., Faccio, R., & Novack, D. (2015). Alternative NF-kB Regulates RANKL-induced Osteoclast Differentiation and Mitochondrial Biogenesis via Independent Mechanisms. J Bone Miner Res, 30(12), 2287–2299.

Zhang, M., Zheng, J., Nussinov, R., & Ma, B. (2017). Release of Cytochrome C from Bax Pores at the Mitochondrial Membrane. Scientific Reports, 7(1).

